# A gastric microbial chromatin remodeler drives gastric cancer progression and immune evasion by reprogramming the host epigenome

**DOI:** 10.64898/2025.12.26.696631

**Authors:** Xiaoshan Xie, Yue Wei, Zhikai Zheng, Jiaying Zheng, Xijie Chen, Jiarui Wang, Ning Ma, Xiaoling Huang, Peng Zhang, Boyu Zhang, Hanyong Cai, Li Ma, Lishi Xiao, Qingxin Liu, Wenyu Wang, Sachiyo Nomura, Shi Chen, Xiangqi Meng, Mong-Hong Lee

**Author notes:** Corresponding author (M.L.), (X.M.), (S.C.). These authors contributed equally to this work (Xiaoshan Xie, Yue Wei).

## Abstract

Emerging evidence highlights the crucial role of microbial communities in modulating cancer development and progression. In gastric cancer (GC), we identify *Streptococcus anginosus* (*SA*) as a tumor-resident oncobacterium that drives both tumor progression and immune evasion. We define a novel mechanism in which *SA* secretes extracellular vesicles (*sa*EVs) that translocate the bacterial chromatin remodeler *sa*SNF2 into host cell. There, *sa*SNF2 partners with host transcription factor TEAD1—through its ATPase activity and participation in BAF complex assembly—to coordinately activate oncogenic transcription. This transkingdom interaction upregulates the palmitoyltransferase ZDHHC11, which in turn stabilizes PD-L1 via palmitoylation to establish an immunosuppressive niche, while also amplifying other TEAD1 target genes to fuel tumor progression. Functionally, *sa*EVs promote tumor growth and limit CD8^+^ T-cell infiltration in vivo. Strikingly, pharmacological inhibition of ZDHHC11 reverses immune evasion and synergizes with anti-PD-1 checkpoint blockade. Our results establish *SA* as a multifaceted driver of GC and reveal the *sa*SNF2-ZDHHC11 axis as a promising target to potentiate immunotherapy.

## Introduction

Gastric cancer (GC) ranks as the fifth most common malignancy and the fifth leading cause of cancer-related death worldwide, accounting for approximately one million new cases and 650,000 deaths annually^1^. Chronic *Helicobacter pylori* (*H. pylori*) infection is the primary known risk factor, yet it progresses to GC in only 1–3% of infected individuals^2^, indicating a critical role for co-factors. Persistent *H. pylori* infection alters gastric physiology and reshapes the resident microbiota, leading to a state of dysbiosis characterized by reduced microbial diversity and enrichment of specific non-*Helicobacter* genera^3, 4, 5, 6^. While such dysbiosis is a hallmark of GC patients, the functional contributions of individual microbes to malignant progression remain largely undefined.

Among emerging bacterial co-factors, *Streptococcus anginosus* (*SA*)—an oral commensal frequently detected in gastric pathologies^7^—has garnered attention. Its abundance often increases as gastric lesions advance, suggesting it may occupy an ecological niche left by receding *H. pylori* levels. Recent work established a direct oncogenic role for *SA*, showing that its surface protein TMPC binds Annexin A2 on gastric epithelial cells to activate the MAPK pathway^4^. However, whether and how *SA* modulates the tumor immune microenvironment (TIME) to facilitate immune evasion and progression is unknown—a gap whose elucidation could reveal new therapeutic avenues.

The tumor immune landscape is a key determinant of both patient outcomes and responses to immune checkpoint inhibitors (ICIs) ^8^. Although combining PD-1 blockade with chemotherapy is now a first-line regimen for advanced GC^9^, most patients still experience primary or acquired resistance. While biomarkers such as programmed cell death ligand 1 (PD-L1) expression, microsatellite instability, and EBV status help guide patient selection^10^, there remains an urgent need to identify novel resistance mechanisms and develop more effective precision immunotherapies^11^.

Here, we identify *SA* as a pivotal gastric oncobacterium that promotes tumor progression by reshaping the cancer cell transcriptome and suppressing antitumor immunity. We uncover a previously unknown cross-kingdom epigenetic mechanism: *SA* releases extracellular vesicles (EVs) that deliver a bacterial chromatin remodeler, *sa*SNF2, into host cells. Within the nucleus, *sa*SNF2 utilizes its ATPase activity to modulate BAF complex and cooperate with the host transcription factor TEAD1, driving the expression of the palmitoyltransferase ZDHHC11. This enzyme in turn stabilizes PD-L1 via palmitoylation, establishing an immunosuppressive TIME and accelerating gastric carcinogenesis. Our findings reveal a microbial-driven axis of immune evasion and propose co-targeting palmitoylation and the PD-1/PD-L1 pathway as a promising strategy for *SA*-positive gastric cancers.

## Results

### *SA* is a tumor-resident oncobacterium that drives gastric carcinogenesis

Analysis of gastric microbiota has consistently shown that gastric cancer (GC) patients harbor a dysbiotic microbial community. Data mining using linear discriminant analysis (LDA) revealed that the relative abundance of *Streptococcus* increases with advancing stages of gastric cancer (Figure 1A). Notably, this increase was more pronounced in metastatic disease, with *Streptococcus* abundance ranking significantly higher in patients with lymph node metastasis (Lp) compared to non-metastatic (Ln) cases (Figure 1B). We validated this spatial association using fluorescence in situ hybridization (FISH), confirming robust *Streptococcus anginosus* (*SA*) enrichment in GC tumor tissues relative to matched adjacent normal tissues (Figure 1C).

**Figure 1.**
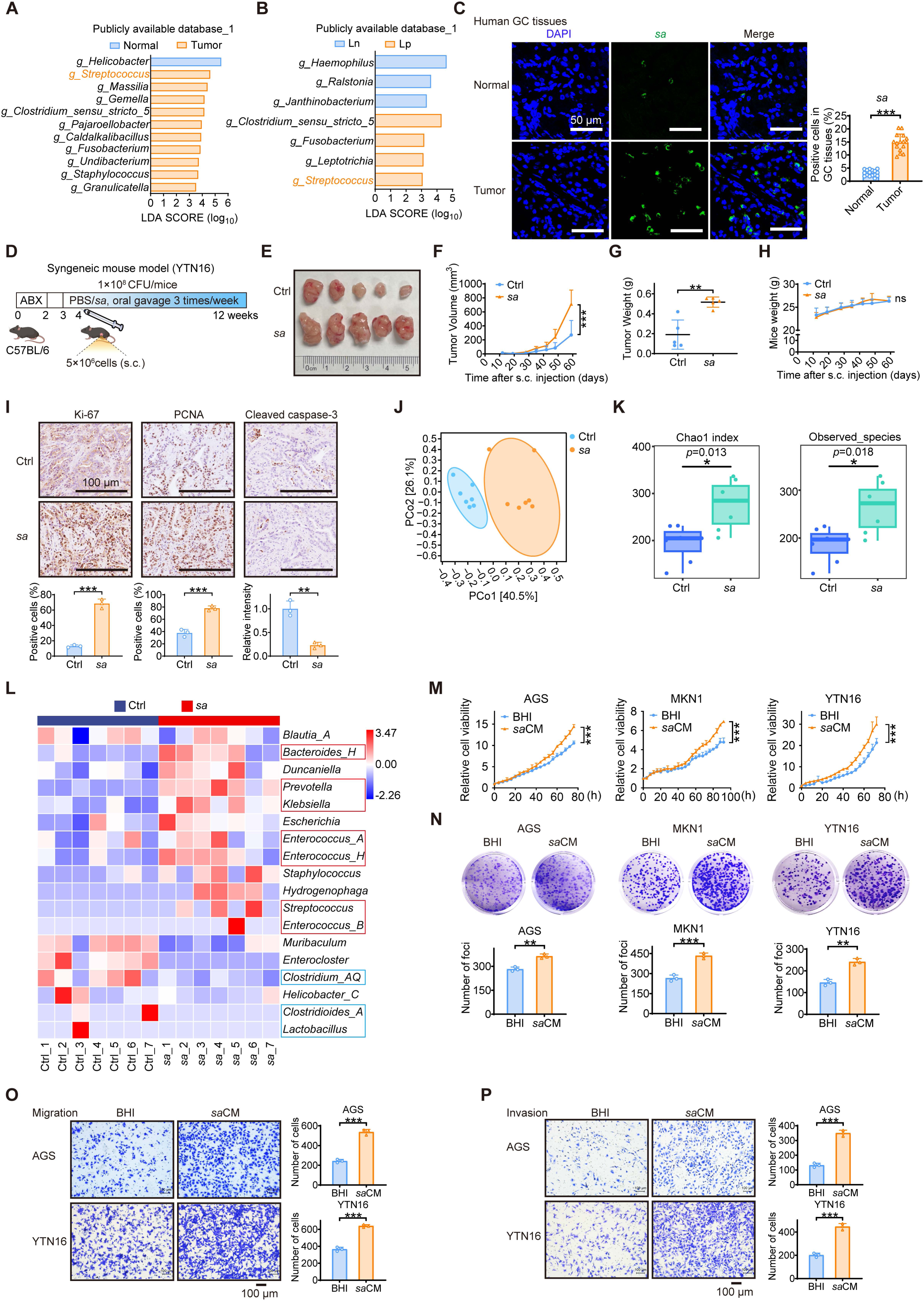
*SA* is a tumor-resident oncobacterium that drives gastric carcinogenesis. (A) Differential bacterial abundance at the genus level in gastric cancer tissues (T) and normal tissues (N), as measured by LDA score. Only taxa with log10 (LDA score) >3 are displayed (n = 40). (B) LDA analysis comparing bacterial abundance in primary gastric cancer lesions between patients with (Lp) and without (Ln) lymph node metastasis. (C) Detection of *SA* in GC tissues by fluorescence in situ hybridization (FISH). Blue (DAPI) indicates nuclei, while green signals represent *SA*-specific probes (left panel). The percentage of *SA*-positive cells in GC tissues was quantified and is shown in a bar graph (right panel). (D) Schematic of the subcutaneous mouse tumor model design. C57BL/6 mice received oral gavage of *SA* (*n* = 5) or PBS (Ctrl, *n* = 5) three times weekly. (E) Illustrative images showing tumors from syngeneic mice (YTN16). (F and G) Data on tumor growth curves and harvested tumor weights are shown. (H) Mice body weights were recorded. (I) Tumor tissue sections were immunohistochemically stained for proliferative markers (Ki67, PCNA) and an apoptosis marker (Cleaved caspase-3) (upper panel). Staining intensity was quantified using ImageJ and presented as bar graphs (lower panel). (J) Principal coordinate analysis (PCoA) based on Bray-Curtis beta diversity of gut microbiota OTUs from mice fecal samples following oral gavage with *SA*. (K) Alpha diversity as measured by the Chao1 index (left panel) and Observed Species (right panel) in mice feces following oral gavage with *SA*. (L) Genus-level heatmap depicting bacterial abundance in mice fecal samples. (M and N) The proliferative capacity and colony formation of AGS, MKN1 and YTN16 cells were examined after treatment with *SA*-conditioned medium (*sa*CM). Control was treated with BHI. (O and P) The migratory and invasive capacities of AGS and YTN16 cells were assessed by Transwell assay after treatment with *sa*CM. Control was treated with BHI. Data are presented as mean ± SD, unpaired Student’s *t* tests. \**p* < 0.05; \*\**p* < 0.01; \*\*\**p* < 0.001; ns, not significant. S.c., subcutaneous; ABX, Antibiotics; OTU, Operational Taxonomic Units; Ctrl, control; BHI, Brain Heart Infusion Broth; CM, conditioned medium.

To directly test the oncogenic role of SA, we established subcutaneous YTN16 gastric cancer tumors in C57BL/6 mice and administered *SA* via oral gavage (Figure 1D). *SA* treatment resulted in a significant increase in both tumor volume (Figures 1E, 1F) and final tumor weight (Figure 1G) compared to the control group, without affecting overall mouse body weight (Figure 1H). Immunohistochemical (IHC) analysis further demonstrated that *SA*-driven tumor growth was associated with elevated expression of the proliferation markers Ki-67 and PCNA and a concurrent reduction in the apoptosis marker cleaved caspase-3 (Figure 1I).

To assess the systemic microbial alterations induced by *SA*, we conducted 16S rRNA sequencing on mouse fecal samples. Principal coordinate analysis (PCoA) revealed distinct clustering, indicating a significant compositional shift in the gut microbiota of *SA*-administered mice compared to the control group (Figure 1J). Consistent with this, alpha diversity analysis using the Chao1 index showed a significant increase in overall microbial richness following *SA* treatment (Figure 1K). Taxonomic profiling

via heatmap analysis indicated that *SA* gavage promoted the relative abundance of several pathobionts, including *Bacteroides*, *Prevotella*, *Klebsiella*, *Enterococcus*, and *Streptococcus* genera (Figure 1L). Concurrently, *SA* administration reduced the abundance of beneficial, homeostatic taxa such as *Clostridium* and *Lactobacillus* (Figure 1L). A combined linear discriminant analysis (LDA) and effect size (LEfSe) analysis (LDA score > 2, *p* < 0.05) confirmed that *SA* treatment induced a marked remodeling of gut microbiota, characterized by the enrichment of pro-tumorigenic communities and the depletion of commensal probiotics (Figures S1A, S1B). Collectively, these data demonstrate that *SA* administration profoundly remodels the gut microbiota into a dysbiotic ecosystem.

To characterize the functional impact of *Streptococcus anginosus* (*SA*) on gastric cancer (GC), we established direct co-culture systems using three GC cell lines (AGS, MKN1, and YTN16). Live *SA* significantly enhanced several oncogenic phenotypes, including proliferation (Figure S1C), colony formation (Figure S1D), migration (Figure S1E), and invasion (Figure S1F). To determine if these effects were mediated by soluble bacterial factors, we treated GC cells with *SA*-conditioned medium (*sa*CM). Remarkably, *sa*CM treatment recapitulated the pro-tumorigenic effects, similarly enhancing cell proliferation (Figure 1M), colony formation (Figure 1N), migration (Figures 1O, S1G), and invasion (Figures 1P, S1H). Together, these results demonstrate that *SA* functions as a potent oncobacterium in GC pathogenesis, and that its tumor-promoting effects are mediated, at least in part, by secreted factors.

### *SA* extracellular vesicles drive tumor progression via dynamin-dependent endocytosis

To identify the key bioactive components in *SA*-conditioned medium (*sa*CM), we performed selective depletion assays. While nuclease treatment (to remove nucleic acids) did not diminish *sa*CM’s pro-proliferative effect, heat inactivation (to denature proteins) abolished it, indicating that proteinaceous components are critical for promoting GC growth (Figures 2A, S2A). Since bacterial extracellular vesicles (EVs) are key mediators of host–microbe communication and efficiently package proteins^12, 13^, we hypothesized that *sa*EVs might be responsible for this activity. We isolated *sa*EVs via ultracentrifugation and confirmed their typical EV morphology by scanning electron microscopy (SEM; Figure 2B) and a size distribution of 50–300 nm (peak 127.6 nm) by nanoparticle tracking analysis (NTA; Figure 2C). Treatment with purified *sa*EVs robustly enhanced GC cell proliferation (Figures 2D, S2B), colony formation (Figures 2E, S2C), migration (Figures 2F, S2D), and invasion (Figures 2G, S2E). Using the lipophilic dye 3,3′-dioctadecyloxacarbocyanine perchlorate (DiO) to label *sa*EVs, we observed efficient internalization by AGS cells within 2 hours via confocal microscopy (Figure 2H). Mechanistically, uptake was blocked by dynasore (a dynamin-dependent endocytosis inhibitor) but not by chlorpromazine (clathrin-mediated endocytosis inhibitor) or cytochalasin D (actin-dependent endocytosis inhibitor), establishing dynamin-dependent endocytosis as the primary entry route for *sa*EVs into GC cells (Figure 2I).

**Figure 2.**
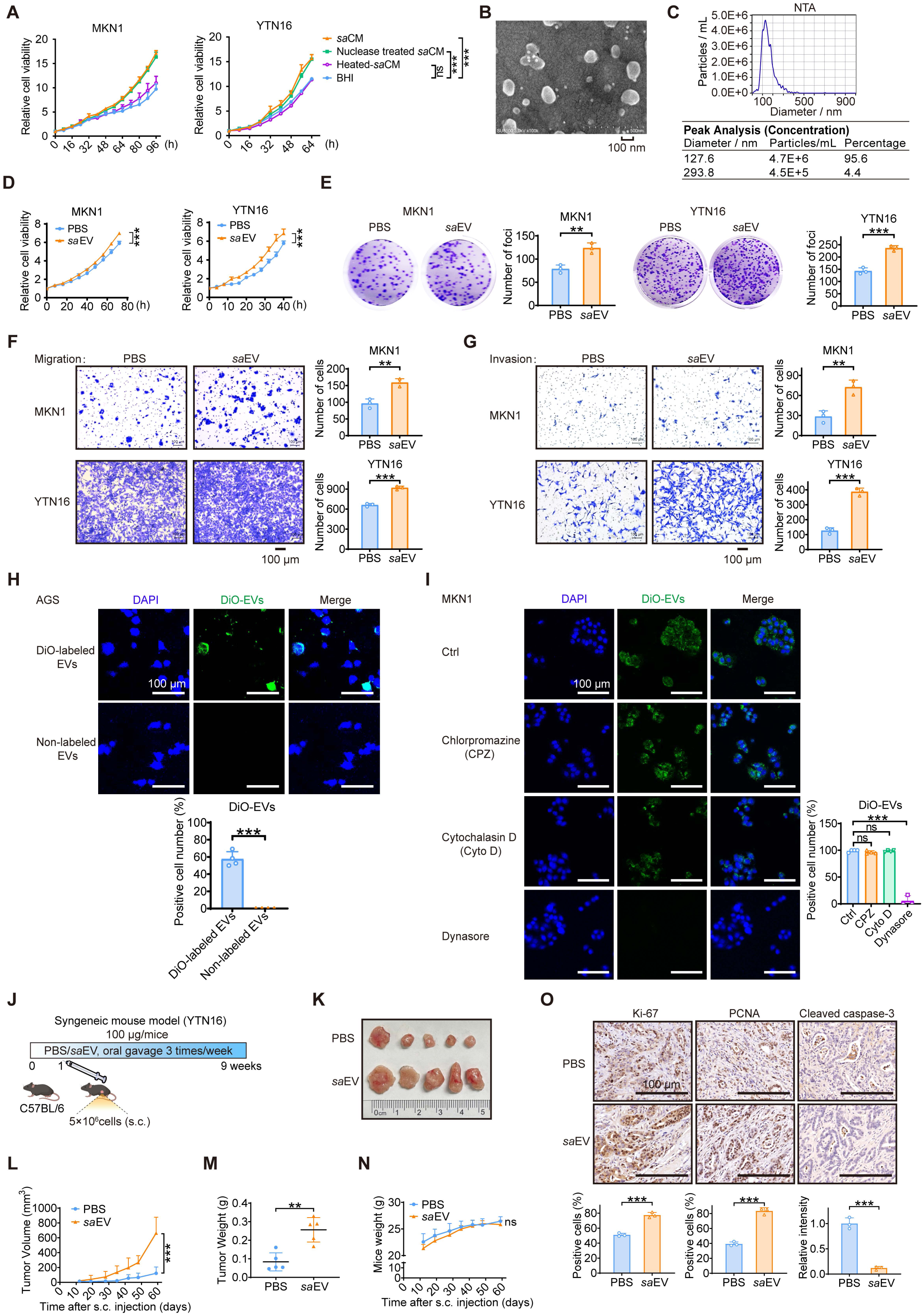
*SA* extracellular vesicles drive tumor progression via dynamin-dependent endocytosis. (A) The proliferative capacity of MKN1 and YTN16 cells were examined after treatment with *sa*CM, nuclease treated *sa*CM or heated *sa*CM. Control was treated with BHI. (B and C) Characterization of *SA*-derived EVs by scanning electron microscopy (SEM) (B) and nanoparticle tracking analysis (NTA) (C). Scale bars, 100 nm. (D and E) The proliferative capacity and colony formation of MKN1 and YTN16 cells were examined after treatment with *sa*EVs. Control was treated with PBS. (F and G) The migratory and invasive capacities of MKN1 and YTN16 cells were assessed by Transwell assay after treatment with *sa*EVs. Control was treated with PBS. (H) Confocal microscopy images demonstrating EVs internalization by AGS cells (20 μg/mL DiO-EVs, 2 h) (upper panel). The number of cells positive for DiO-labeled EVs was quantified and is shown in a bar graph (lower panel). Scale bars, 100 μm. (I) Confocal images of MKN1 cells pretreated with specific endocytosis inhibitors (1 h) prior to 2-hour exposure to DiO-labeled *sa*EVs (20 μg/mL) (left panel). The number of DiO-EVs-positive cells was quantified and is shown in a bar graph (right panel). Scale bars, 100 μm. (J) Schematic of the subcutaneous mouse model design. C57BL/6 mice received oral gavage of *sa*EVs (*n* = 5) or PBS (*n* = 5) three times weekly. (K) Illustrative images showing tumors from syngeneic mice (YTN16). (L and M) Data on tumor growth curves and harvested tumor weights are shown. (N) Mice body weights were recorded. (O) Tumor tissue sections were immunohistochemically stained for proliferative markers (Ki67, PCNA) and an apoptosis marker (Cleaved caspase-3) (upper panel). Staining intensity was quantified using ImageJ and presented as bar graphs (lower panel). Data are presented as mean ± SD. \**p* < 0.05; \*\**p* < 0.01; \*\*\**p* < 0.001; ns, not significant. BHI, Brain Heart Infusion Broth; CM, conditioned medium; EV, extracellular vesicle; s.c., subcutaneous.

In a subcutaneous syngeneic mouse model using YTN16 GC cells, oral gavage of *sa*EVs significantly increased tumor volume (Figures 2J–2K) and final tumor weight (Figure 2M), with no adverse effect on overall mouse body weight (Figure 2N). Immunohistochemical analysis of these tumors revealed a concomitant increase in the proliferation markers Ki-67 and PCNA and a decrease in the apoptosis marker cleaved caspase-3 (Figure 2O). Taken together, these results demonstrate that *sa*EVs are internalized by GC cells via dynamin-dependent endocytosis and that they function as critical drivers of malignancy, enhancing tumor cell proliferation and accelerating tumor growth in vivo.

### *sa*EVs drive gastric cancer immune evasion by upregulating the palmitoyltransferase ZDHHC11 to stabilize PD-L1

To define the molecular mechanism by which *sa*EVs promote gastric cancer (GC) progression, we performed RNA sequencing on AGS cells following *sa*EV or PBS (control) treatment. Gene Ontology (GO) enrichment analysis revealed a pronounced enrichment of pathways associated with protein palmitoylation, palmitoyltransferase activity, and peptidyl-cysteine modification in *sa*EV-treated cells (Figure 3A). Volcano plot analysis of differentially expressed genes identified several candidates within these pathways, with the palmitoyltransferase *ZDHHC11* showing the most significant upregulation (Figure 3B). We confirmed this finding using qRT-PCR, demonstrating consistent *ZDHHC11* induction by *sa*EVs across multiple GC cell lines—AGS, MKN1, MKN45, and YTN16 (Figure 3C).

**Figure 3.**
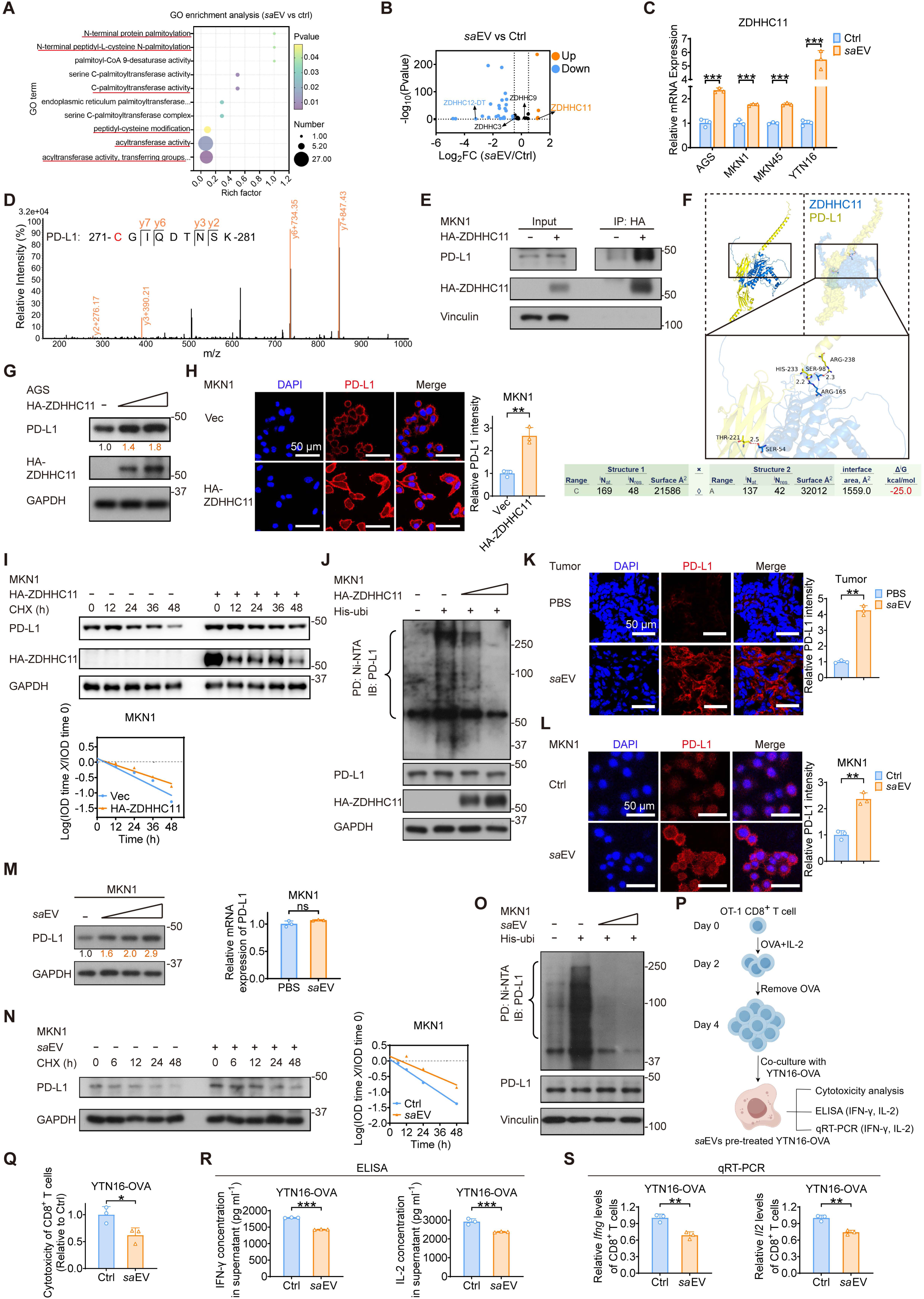
*sa*EVs drive gastric cancer immune evasion by upregulating the palmitoyltransferase ZDHHC11 to stabilize PD-L1. (A) Gene Ontology (GO) enrichment analysis of pathways enriched in the *sa*EV treatment group in AGS cells, as identified by RNA sequencing. (B) Volcano plot of differentially expressed genes in the pathway from (A) after *sa*EV treatment. (C) qRT-PCR analysis demonstrating an upregulation of ZDHHC11 in GC cells following treatment with *sa*EV. (D) Peak map of PD-L1 from immunoprecipitation mass spectrometry. HA-tagged ZDHHC11 plasmid was transiently transfected into AGS cells; HA antibody and Protein A/G beads were used to pull down ZDHHC11-interacting proteins. (E) MKN1 cells were transfected with HA-tagged ZDHHC11, immunoprecipitated using HA antibody with Protein A/G beads, and immunoblotted with indicated antibodies. (F) Three-dimensional (3D) structural representation of potential binding sites between ZDHHC11 and PD-L1, with hydrogen bonds indicated (upper panel). The calculated binding free energy was -25.0 kcal/mol (lower panel). (G) Immunoblot analysis of PD-L1 upregulation in AGS cells following ZDHHC11 overexpression. (H) Immunofluorescence staining demonstrating PD-L1 upregulation in MKN1 cells after ZDHHC11 overexpression (left panel). PD-L1 fluorescence intensity was quantified using ImageJ (right panel). (I) Following ZDHHC11 overexpression, MKN1 cells were treated with cycloheximide (CHX; 100 μg/mL) for specified durations. PD-L1 protein stability was assessed by immunoblotting and quantified using ImageJ. (J) MKN1 cells carrying the indicated plasmids were harvested following 6-hour treatment with 25 μM MG132. Cell lysates were subjected to nickel beads pulldown assays and immunoblotted with indicated antibodies. (K) Immunofluorescence staining showing PD-L1 upregulation in mice tumor tissues following oral administration of *sa*EVs (left panel). PD-L1 fluorescence intensity was quantified using ImageJ (right panel). (L) Immunofluorescence staining demonstrating PD-L1 upregulation in MKN1 cells after *sa*EV treatment (left panel). PD-L1 fluorescence intensity was quantified using ImageJ (right panel). (M) Immunoblot analysis showing PD-L1 protein upregulation after *sa*EV treatment (left panel). qRT-PCR analysis reveals unchanged PD-L1 mRNA levels in *sa*EV-treated MKN1 cells (right panel). (N) Following *sa*EV treatment, MKN1 cells were treated with cycloheximide (CHX; 100 μg/mL) for specified durations. PD-L1 protein stability was assessed by immunoblotting and quantified using ImageJ. (O) MKN1 cells treated with *sa*EV were harvested following 6-hour exposure to 25 μM MG132. Cell lysates were subjected to nickel beads pulldown assays and immunoblotted with indicated antibodies. (P) Schematic diagram of the coculture process between YTN16-OVA cells (stably expressing ovalbumin) and CD8^+^ T cells. (Q to S) YTN16-OVA cells were pretreated with *sa*EVs before 24-hour coculture with activated splenic CD8^+^ T cells from OT-1 mice. Cytotoxicity of CD8^+^ T cells against YTN16-OVA cells was measured (Q). Supernatant concentrations of IL-2 and IFN-γ were determined by ELISA (R). *Il2* and *Ifng* mRNA expression levels in CD8^+^ T cells were analyzed by qRT-PCR (S). Data are presented as mean ± SD. \**p* < 0.05; \*\**p* < 0.01; \*\*\**p* < 0.001; ns, not significant. EV, extracellular vesicle; CHX, cycloheximide; PD, pulldown; IB, immunoblot; OVA, ovalbumin.

To identify ZDHHC11 substrates, we performed immunoprecipitation-mass spectrometry (IP-MS), which revealed a significant enrichment of PD-L1-derived peptides in ZDHHC11 immunoprecipitates compared to controls (Figure 3D), indicating a direct interaction. Subsequent co-immunoprecipitation (co-IP) assays confirmed this physical association (Figure 3E). Molecular docking further predicted a high-affinity interaction, with a binding free energy of –25 kcal/mol (Figure 3F). While other palmitoyltransferases (ZDHHC3 and ZDHHC9) have been reported to modify PD-L1^14, 15^, qRT-PCR showed that *sa*EVs specifically upregulate *ZDHHC11* without altering *ZDHHC3* or *ZDHHC9* expression (Figure S3A), indicating pathway selectivity.

Functionally, overexpressing ZDHHC11 increased PD-L1 steady-state protein levels in a dose-dependent manner, as shown by immunoblotting (Figures 3G, S3B) and immunofluorescence intensity (Figure 3H). ZDHHC11 overexpression also significantly decelerated PD-L1 turnover in MKN1 and MKN45 cells (Figures 3I, S3C) and reduced its polyubiquitination in a do se-dependent manner (Figure 3J). Conversely, doxycycline-induced ZDHHC11 knockdown increased PD-L1 polyubiquitination in AGS and MKN45 cells (Figure S3D). Together, these results establish ZDHHC11 as a PD-L1-specific palmitoyltransferase that stabilizes PD-L1 by inhibiting its polyubiquitin-mediated degradation.

To determine whether *sa*EVs modulate PD-L1 expression in vivo via ZDHHC11 upregulation, we analyzed tumor tissues from our murine models. Immunofluorescence (IF) staining revealed that *sa*EV administration significantly increased PD-L1 protein levels in tumors (Figures 3K, S3E), a finding corroborated in *sa*EV-treated MKN1 cells in vitro (Figure 3L). Notably, this elevation occurred post-transcriptionally, as PD-L1 mRNA levels were unaffected (Figures 3M, S3F). Consistent with a post-translational mechanism, cycloheximide (CHX) chase assays demonstrated that *sa*EVs decelerated PD-L1 turnover in MKN1 and MKN45 cells (Figures 3N, S3G). Furthermore, *sa*EV treatment reduced PD-L1 polyubiquitination (Figures 3O, S3H), confirming its role in stabilizing PD-L1.

Given PD-L1’s immunosuppressive function, we next assessed the functional consequence of *sa*EV-mediated stabilization using an OT-1 CD8^+^ T cell coculture assay (Figure 3P). GC cells pretreated with *sa*EVs significantly impaired CD8^+^ T cell cytotoxicity (Figure 3Q) and suppressed the secretion of effector cytokines IFN-γ and IL-2, as measured by ELISA and qRT-PCR (Figures 3R, 3S). Collectively, these results establish that *sa*EVs upregulate ZDHHC11 to stabilize PD-L1, which in turn suppresses CD8^+^ T cell effector function and promotes tumor immune evasion.

### *sa*EVs stabilize PD-L1 and potentiate immune evasion via ZDHHC11-mediated palmitoylation at Cys272

To directly test whether *sa*EVs regulate PD-L1 stability via palmitoylation, we performed site-specific analysis. Liquid chromatography-mass spectrometry (LC-MS) identified palmitoylation at the Cys272 residue of PD-L1 and showed that *sa*EV treatment significantly increased its normalized palmitoylation levels compared to control (Figures 4A, 4B). We validated this finding using an acyl-biotin exchange (ABE) assay, which confirmed that *sa*EVs elevate PD-L1 palmitoylation—an effect that was reversible upon treatment with the palmitoylation inhibitor 2-bromopalmitate (2-BP) (Figure 4C).

**Figure 4.**
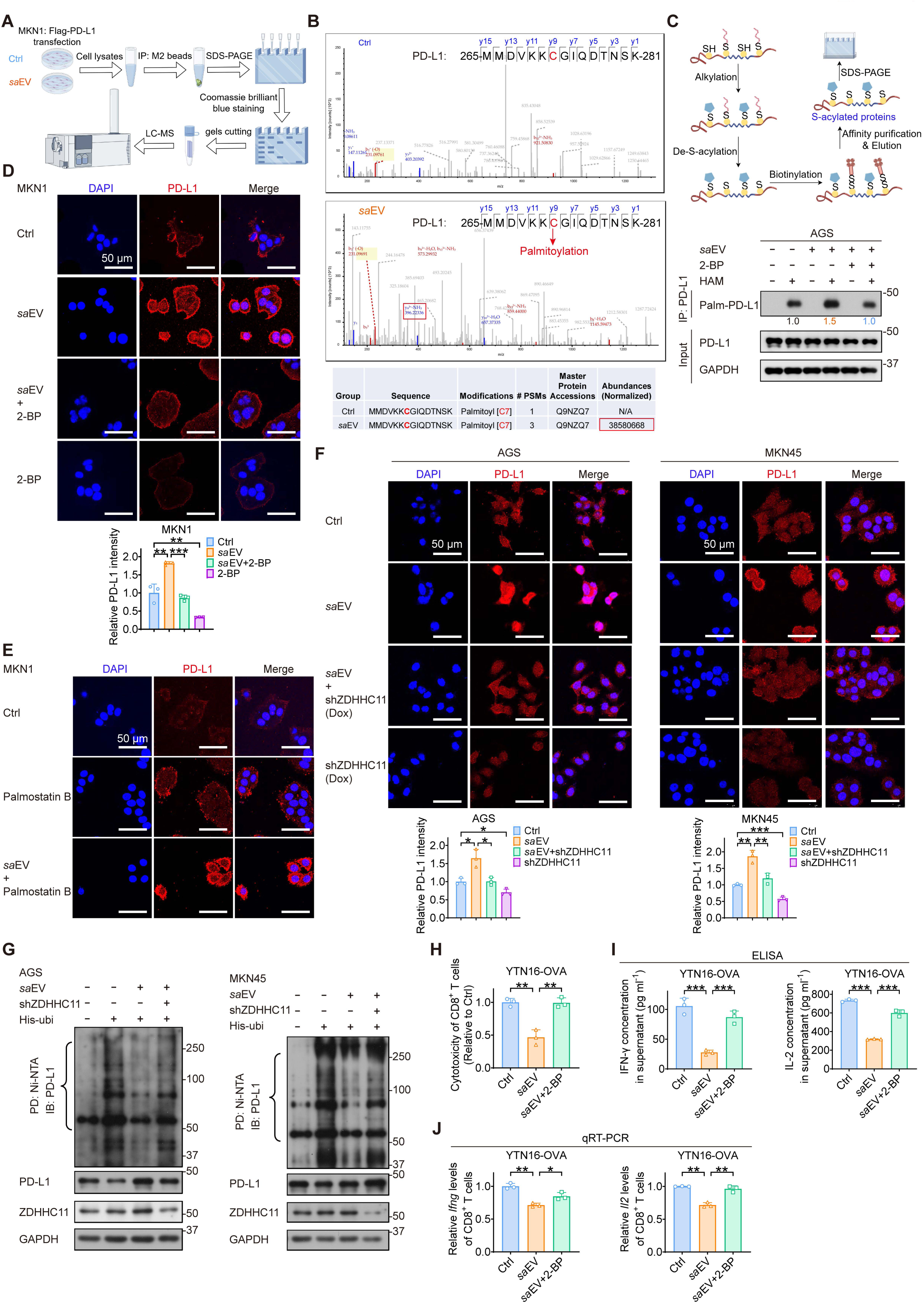
*sa*EVs stabilize PD-L1 and potentiate immune evasion via ZDHHC11-mediated palmitoylation at Cys272. (A) Schematic diagram of the LC-MS process for PD-L1 modification analysis in MKN1 cells treated with *sa*EVs or PBS (control). MKN1 cells transiently transfected with Flag-tagged PD-L1 plasmids were treated with *sa*EVs or PBS. PD-L1 was immunoprecipitated using anti-Flag M2 beads, and its palmitoylation status was analyzed by LC-MS. (B) Peak maps of PD-L1 from LC-MS after treatment with *sa*EVs or PBS (control). (C) Schematic diagram of the ABE assay workflow for palmitoylation detection (upper panel). Detection of palmitoylated PD-L1 (Palm-PD-L1) in Flag-PD-L1-expressing AGS cells treated with either *sa*EVs (50 μg/mL, 24 h) or 2-BP (50 μM, 24 h), using the IP-ABE assay (lower panel). (D) Immunofluorescence staining showing PD-L1 upregulation in MKN1 cells after treatment with *sa*EVs (50 μg/mL, 24 h), which was reversed by 2-BP (50 μM, 24 h) (upper panel). PD-L1 fluorescence intensity was quantified using ImageJ (lower panel). (E) Immunofluorescence staining showing PD-L1 upregulation in MKN1 cells after treatment with *sa*EVs (50 μg/mL, 24 h) and palmostatin B (5 μM, 24 h) treatment. (F) Immunofluorescence staining showing PD-L1 upregulation in AGS and MKN45 cells after treatment with *sa*EVs (50 μg/mL, 24 h), which was reversed upon ZDHHC11 knockdown (Dox-incucible) (upper panel). PD-L1 fluorescence intensity was quantified using ImageJ (lower panel). (G) AGS and MKN45 cells carrying the indicated plasmids were harvested following 6-hour treatment with 25 μM MG132. Cell lysates were subjected to nickel beads pulldown assays and immunoblotted with indicated antibodies. (H to J) YTN16-OVA cells were pretreated with *sa*EVs and 2-BP before 24-hour coculture with activated splenic CD8^+^ T cells from OT-1 mice. Cytotoxicity of CD8^+^ T cells against YTN16-OVA cells was measured (H). Supernatant concentrations of IL-2 and IFN-γ were determined by ELISA (I). *Il2* and *Ifng* mRNA expression levels in CD8^+^ T cells were analyzed by qRT-PCR (J). Data are presented as mean ± SD. \**p* < 0.05; \*\**p* < 0.01; \*\*\**p* < 0.001; ns, not significant. LC-MS, Liquid chromatography–mass spectrometry; ABE, Acyl-biotin exchange; 2-BP, 2-Bromohexadecanoic acid.

Immunofluorescence staining confirmed that *sa*EV-induced PD-L1 upregulation was suppressed by 2-BP (Figure 4D), while combined treatment with the palmitoylation inducer palmostatin B further enhanced PD-L1 expression (Figure 4E). Furthermore, doxycycline-induced ZDHHC11 knockdown in AGS and MKN45 cells attenuated the *sa*EV-mediated increase in PD-L1 (Figure 4F), underscoring the critical role of ZDHHC11 in this process. Consistent with this, ZDHHC11 knockdown also reversed the *sa*EV-mediated attenuation of PD-L1 polyubiquitination (Figure 4G), indicating that this phenomenon is ZDHHC11-dependent.

Functional assays using the OT-1 coculture system demonstrated that pretreatment of YTN16 cells with *sa*EVs significantly suppressed CD8^+^ T cell cytotoxicity—an effect that was reversed by 2-BP treatment (Figure 4H). Consistent with this, *sa*EVs inhibited IFN-γ and IL-2 production, as measured by ELISA and qRT-PCR; these effects were also reversed by 2-BP administration (Figures 4I and 4J). Collectively, these data demonstrate that *sa*EVs enhance PD-L1 stability via ZDHHC11-mediated palmitoylation, thereby promoting immune evasion in the GC microenvironment.

### The *sa*EV-derived chromatin remodeler *sa*SNF2 enhances TEAD1-mediated transcription of the ZDHHC11 gene

To investigate how *sa*EVs upregulate ZDHHC11, we first used the JASPAR database to predict transcription factors that might regulate its transcription. This analysis identified TEAD1, MEF2A, and IRF1 as top candidates based on binding affinity scores (Figure 5A). Functional validation in gastric cancer cell lines (MKN1 and AGS) showed that TEAD1 overexpression significantly upregulated ZDHHC11 mRNA in a dose-dependent manner (Figures 5B, S4B), whereas MEF2A and IRF1 had no such effect (Figure S4A). Pan-cancer analysis via GEPIA indicated elevated TEAD1 expression in stomach adenocarcinoma (STAD) compared to normal tissues (Figure S4C). Notably, high expression of either TEAD1 or ZDHHC11 was associated with poor survival in gastrointestinal cancers (STAD, COAD, and READ) based on TCGA data (Figures S4D, S4E). Additionally, TEAD1 expression positively correlated with ZDHHC11 across multiple gastrointestinal cancers (STAD, COAD, READ, ESCA, and LIHC) in the TCGA dataset (Figure S4F). Chromatin immunoprecipitation (ChIP) assays confirmed TEAD1 binding to five putative sites (S1–S5) within the ZDHHC11 promoter (Figures 5C, 5D), indicating that TEAD1 directly regulates ZDHHC11 transcription. However, *sa*EV administration did not alter TEAD1 protein levels, as shown by IHC staining (Figure S4G). We therefore hypothesize that *sa*EV-derived factors may activate TEAD1, thereby enhancing ZDHHC11 expression.

**Figure 5.**
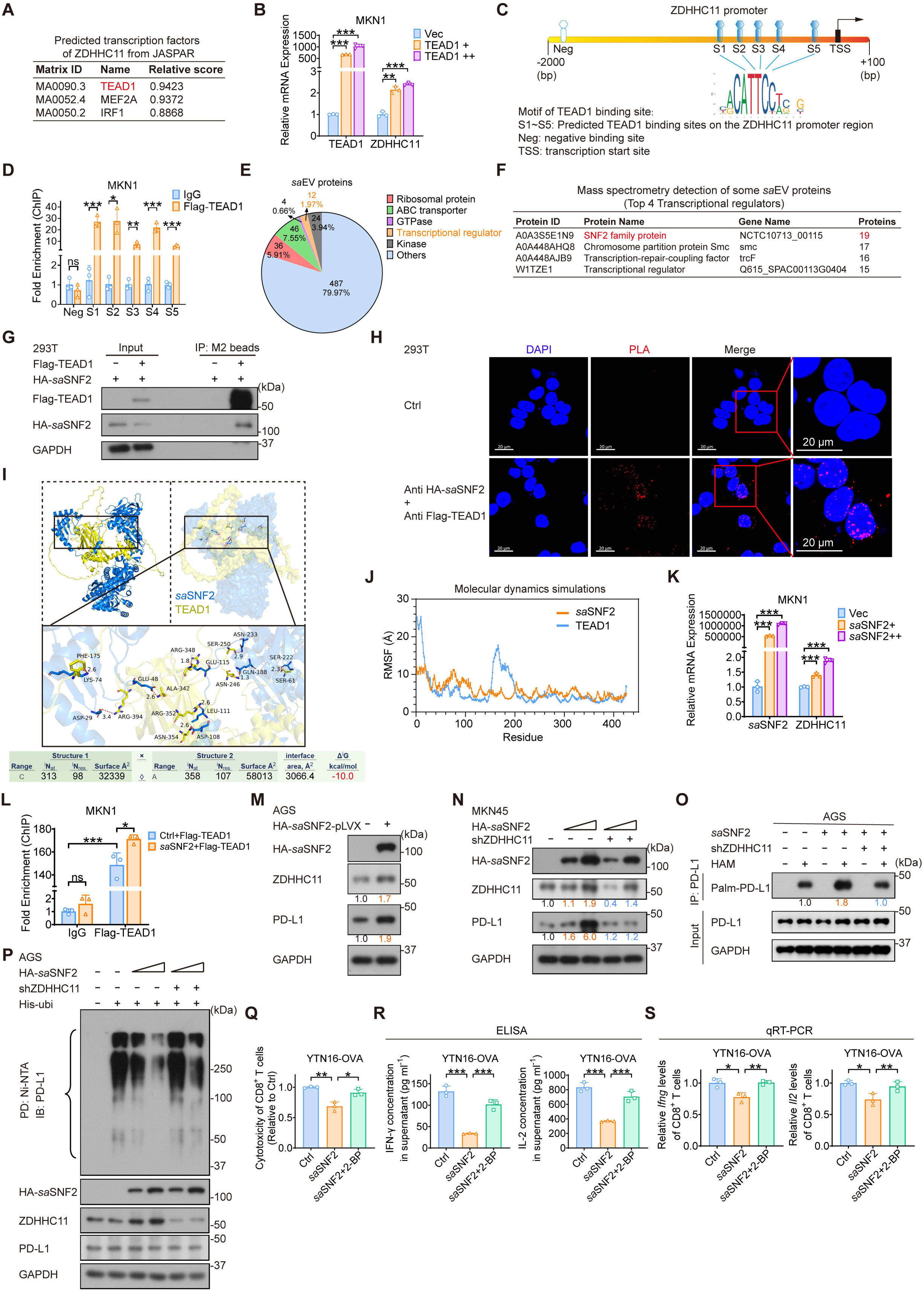
The *sa*EV-derived chromatin remodeler *sa*SNF2 enhances TEAD1-mediated transcription of the ZDHHC11 gene. (A) Predicted transcription factors of ZDHHC11 from JASPAR website. (B) qRT-PCR analysis demonstrating an upregulation of ZDHHC11 in MKN1 cells following TEAD1 overexpression. (C) Five potential TEAD1-binding sites (S1–S5) were predicted within the ZDHHC11 promoter region (−2,000 to +100 bp). (D) Chromatin immunoprecipitation (ChIP) assays in MKN1 cells validated TEAD1 occupancy at multiple distinct sites within the ZDHHC11 promoter region. (E) Proteins identified in *sa*EVs by 4D label-free quantitative proteomics. (F) The Top 4 transcriptional regulator proteins identified in *sa*EVs. (G) 293T cells were transfected with Flag-tagged TEAD1 and HA-tagged *sa*SNF2, immunoprecipitated with M2 beads, and immunoblotted with indicated antibodies. (H) Proximity ligation assay (PLA) was performed in 293T cells co-transfected with Flag-tagged TEAD1 and HA-tagged *sa*SNF2. HA/Flag antibody-based detection revealed proximal interactions (red signals), with nuclear counterstaining by DAPI (blue). (I) Three-dimensional (3D) structural representation of potential binding sites between *sa*SNF2 and TEAD1, with hydrogen bonds indicated. The calculated binding free energy was −10.0 kcal/mol. (J) Molecular dynamics simulations revealed the root-mean-square fluctuation (RMSF) values of *sa*SNF2 (orange) and TEAD1 (blue). (K) qRT-PCR analysis demonstrating an upregulation of ZDHHC11 in MKN1 cells following *sa*SNF2 overexpression. (L) *sa*SNF2 overexpression enhanced TEAD1 recruitment to the ZDHHC11 promoter, as quantified by ChIP assay. (M and N) Immunoblot analysis of ZDHHC11 and PD-L1 expression in AGS and MKN45 cells following *sa*SNF2 overexpression and ZDHHC11 knockdown (Dox-inducible). (O) Detection of palmitoylated PD-L1 (Palm-PD-L1) in Flag-PD-L1-expressing AGS cells under the indicated treatment, using the IP-ABE assay. (P) AGS cells carrying the indicated plasmids were harvested following 6-hour treatment with 25 μM MG132. Cell lysates were subjected to nickel beads pulldown assays and immunoblotted with indicated antibodies. (Q to S) YTN16-OVA cells were transfected with *sa*SNF2 overexpression plasmids and pretreated with 2-BP before 24-hour coculture with activated splenic CD8^+^ T cells from OT-1 mice. Cytotoxicity of CD8^+^ T cells against YTN16-OVA cells was measured (Q). Supernatant concentrations of IL-2 and IFN-γ were determined by ELISA (R). *Il2* and *Ifng* mRNA expression levels in CD8^+^ T cells were analyzed by qRT-PCR (S). Data are presented as mean ± SD. \**p* < 0.05; \*\**p* < 0.01; \*\*\**p* < 0.001; ns, not significant. ChIP, Chromatin immunoprecipitation; EV, extracellular vesicle; PLA, Proximity ligation assay; RMSF, root-mean-square fluctuation; ABE, Acyl-biotin exchange; 2-BP, 2-Bromohexadecanoic acid; PD, pulldown; IB, immunoblot.

To identify *sa*EV factors that activate TEAD1, we performed proteomic analysis via mass spectrometry and identified 12 proteins potentially involved in TEAD1-mediated transcription (Figure 5E). The top candidate, *SA*-SNF2 (hereafter *sa*SNF2; SNF: SUCROSE NONFERMENTING), a helicase-like chromatin remodeler, was selected for further investigation (Figures 5E, 5F). Structural alignment revealed 55% similarity between *sa*SNF2 and human Brahma-related gene 1 (BRG1) within their ATP-binding domains (Figure S4H). Co-immunoprecipitation (Co-IP) and proximity ligation assays (PLA) confirmed a physical interaction between *sa*SNF2 and TEAD1 (Figures 5G, 5H). Molecular docking and dynamics simulations supported a stable *sa*SNF2-TEAD1 complex, with a binding energy of −10 kcal/mol, a low RMSF (<10 Å), and an equilibrated RMSD of ∼11.2 Å (Figures 5I, 5J, and S4I). Functionally, *sa*SNF2 overexpression increased ZDHHC11 mRNA in GC cells (MKN1, AGS, YTN16) in a dose-dependent manner (Figures 5K, S4J) and enhanced TEAD1 recruitment to the ZDHHC11 promoter (Figure 5L). Accordingly, *sa*SNF2 upregulated ZDHHC11, leading to elevated PD-L1 protein levels (Figures 5M, S4K), whereas ZDHHC11 knockdown attenuated this *sa*SNF2-mediated PD-L1 elevation (Figures 5N, S4L).

Acyl-biotin exchange (ABE) assays confirmed that the *sa*SNF2-mediated enhancement of PD-L1 palmitoylation depends on ZDHHC11, as ZDHHC11 knockdown abrogated this effect (Figure 5O). Furthermore, *sa*SNF2 reduced PD-L1 ubiquitination in a ZDHHC11-dependent manner (Figures 5P, S4M). In an OT-1 coculture system, the *sa*SNF2-driven immune evasion—characterized by diminished CD8^+^ T-cell cytotoxicity and IFN-γ/IL-2 production—was reversed by the palmitoylation inhibitor 2-BP (Figures 5Q–5S). Collectively, these data establish that *sa*SNF2 activates TEAD1 to transactivate ZDHHC11, thereby promoting PD-L1 palmitoylation and immune evasion in gastric cancer.

### The ATPase motif of *sa*SNF2 is critical for TEAD1-mediated transcription and for assembly of the host BAF complex

Sequence similarity analysis of full-length *sa*SNF2 across bacterial species and human BRG1 (Brahma-related gene 1) revealed low interspecies similarity (<40%) (Figure S5A). Despite this overall divergence, a highly conserved ATPase motif (DXMGLGKT) was identified (Figure 6A), suggesting conservation of ATPase function. To investigate the functional role of this motif, we generated a mutant (*sa*SNF2-A8 mut) in which the motif was substituted with alanine (AAAAAAAA) (Figure 6A). Structural modeling in PyMOL highlighted the 3D conformation of the wild-type and mutant ATPase motifs (Figure 6B). Functional assays showed that while overexpression of wild-type *sa*SNF2 (*sa*SNF2-WT) enhanced ZDHHC11 mRNA expression in MKN1, AGS, and YTN16 cells, the *sa*SNF2-A8 mutant lost this capacity (Figures 6C, S5B).

**Figure 6.**
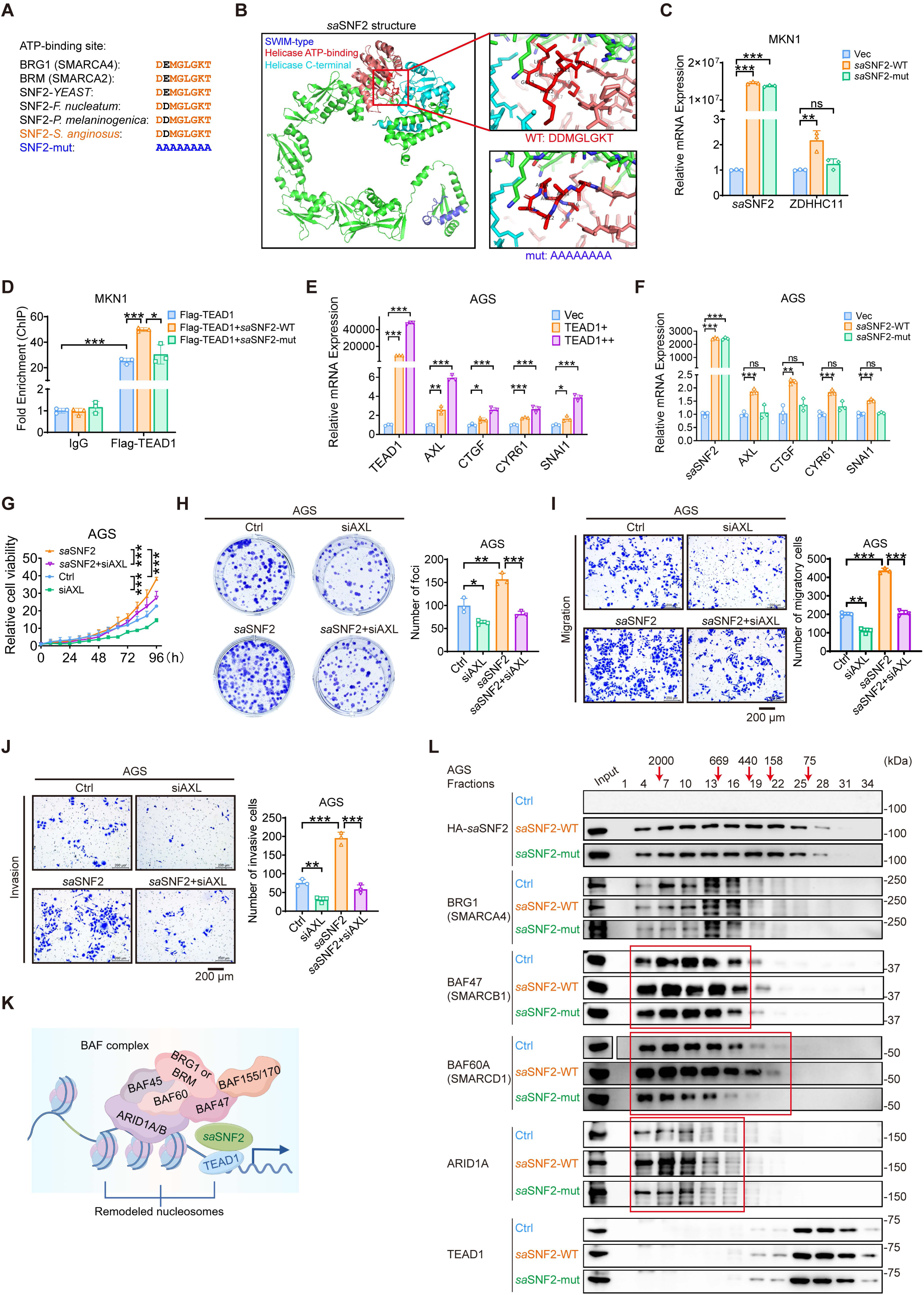
The ATPase motif of *sa*SNF2 is critical for TEAD1-mediated transcription and for assembly of the host BAF complex. (A) ATP-binding site and its mutation variants in different species. (B) A schematic diagram of *sa*SNF2’s secondary structure (three domains) and its ATP-binding site (wild-type: DDMGLGKT; mutant: AAAAAAAA) was visualized using PyMOL. (C) qRT-PCR analysis demonstrating ZDHHC11 expression in MKN1 cells following overexpression of wild-type or mutant *sa*SNF2. (D) ChIP assays showing TEAD1 binding to the ZDHHC11 promoter region in MKN1 cells after overexpression of wild-type or mutant *sa*SNF2. (E) qRT-PCR analysis demonstrating TEAD1 target genes expression in AGS cells following overexpression of TEAD1. (F) qRT-PCR analysis demonstrating TEAD1 target genes expression in AGS cells following overexpression of wild-type or mutant *sa*SNF2. (G and H) The proliferative capacity and colony formation of AGS cells were examined after *sa*SNF2 overexpression and AXL knockdown (siRNA). (I and J) The migratory and invasive capacities of AGS cells were assessed by Transwell assay after *sa*SNF2 overexpression and AXL knockdown (siRNA). (K) Schematic diagram of the human BAF (BRG1/BRM-associated factor) chromatin-remodeling complex. (L) Gel filtration chromatography analysis of AGS cell lysates expressing empty vector (Ctrl), wild-type *sa*SNF2 (*sa*SNF2-WT), or mutant *sa*SNF2 (*sa*SNF2-mut). The eluted fraction’s molecular weight is indicated above. Data are presented as mean ± SD. \**p* < 0.05; \*\**p* < 0.01; \*\*\**p* < 0.001; ns, not significant. ChIP, Chromatin immunoprecipitation; BAF, BRG1/BRM-associated factor.

Moreover, ChIP assays revealed that the *sa*SNF2-A8 mutant failed to facilitate TEAD1 binding to the ZDHHC11 promoter, in contrast to the wild-type protein (Figure 6D), indicating that the ATPase activity is critical for this recruitment. Given the established role of TEAD transcription factors in activating pro-tumorigenic genes such as *AXL*, *CTGF*, *CYR61*, and *SNAI1*^16, 17^, we investigated whether *sa*SNF2 modulates these targets. Indeed, TEAD1 overexpression upregulated AXL, CTGF, CYR61 and SNAI1 in AGS cells (Figure 6E), a finding corroborated by positive correlations in the GEPIA database (Figure S5C). Intriguingly, *sa*SNF2-WT, but not *sa*SNF2-A8 mut, similarly activated these genes (Figure 6F). AXL, a member of the TAM (TYRO3, AXL, MER) receptor tyrosine kinase family and a transforming gene in cells from chronic myeloproliferative disorder patients, is a TEAD target and proliferation promoting gene overexpressed in various types of cancer^18, 19^. Importantly, *sa*SNF2-driven oncogenic phenotypes (growth, colony formation, migration, invasion) were reversed upon AXL silencing (Figures 6G–6J, S5D–S5G), suggesting a *sa*SNF2-TEAD-AXL link. Collectively, the data demonstrate that the ATPase motif of *sa*SNF2 is essential for TEAD1-dependent transcription, governing both *ZDHHC11* and key pro-tumorigenic target genes.

The BRG1/BRM-associated factor (BAF) complex is a multi-subunit chromatin remodeling machinery that is central to transcriptional regulation and is shared among all three subfamilies of the SWI/SNF (switching defective/sucrose nonfermenting) complexes (Figure 6K). This complex functions by using ATP hydrolysis to reposition nucleosomes, eject histone octamers, or evict histone dimers^20^. The catalytic core of the BAF complex is BRG1, an ATPase motor subunit that translocates DNA along the nucleosome^20, 21^. Given that *sa*SNF2 shares homology with BRG1 and possesses conserved ATPase activity, we hypothesized that it might incorporate into or functionally engage with the host BAF complex to modulate its chromatin remodeling activity.

Gel filtration assays showed that *sa*SNF2-WT altered BAF complex composition, while *sa*SNF2-mut did not (Figure 6L), indicating that *sa*SNF2 ATPase activity contributes to BAF complex assembly. Specifically, the presence of *sa*SNF2 increased the distribution of ARID1A—the second most frequently mutated gene in GC^22^—across BAF complexes (Figure 6L). Similarly, the distribution of BAF60A, which links transcription factors to the BAF complex to regulate gene expression^23^, and BAF47, whose basic α helix binds the nucleosomal acidic patch^24^, also increased with *sa*SNF2. In contrast, BRG1 distribution across BAF complexes remained largely unaffected (Figure 6L). Although TEAD1 partially associates with the BAF complex, its distribution was not substantially altered by *sa*SNF2 (Figure 6L).

SWI/SNF subunits assemble into diverse chromatin-remodeling complexes that vary by cell type and developmental stage^23^, with humans capable of forming over 1,400 distinct SWI/SNF complexes across tissues^25^. Thus, *sa*SNF2 appears to participate in the dynamic assembly of BAF complexes, potentially facilitating the function of specialized complexes in regulating transcription of certain target genes, such as *TEAD1*, through epigenetic mechanisms.

### *sa*SNF2 gene expression in tumor cells recapitulates the *SA*-mediated tumor growth promotion and PD-L1 palmitoylation/stabilization by upregulating ZDHHC11

To investigate the oncogenic role of *sa*SNF2 in vivo, we established a syngeneic mouse model by subcutaneously injecting C57BL/6 mice with YTN16 cells stably overexpressing pLVX-*sa*SNF2-HA (Figure 7A). Compared with controls, *sa*SNF2 overexpression significantly increased both tumor volume (Figures 7B, 7C) and tumor weight (Figure 7D). Immunohistochemical (IHC) analysis further showed elevated Ki-67 and decreased cleaved caspase-3 in *sa*SNF2-overexpressing tumors, along with upregulated expression of ZDHHC11 and PD-L1 (Figure 7E). These results demonstrate that *sa*SNF2 alone is sufficient to promote gastric cancer (GC) progression in vivo, an effect mediated through activation of the ZDHHC11–PD-L1 axis.

**Figure 7.**
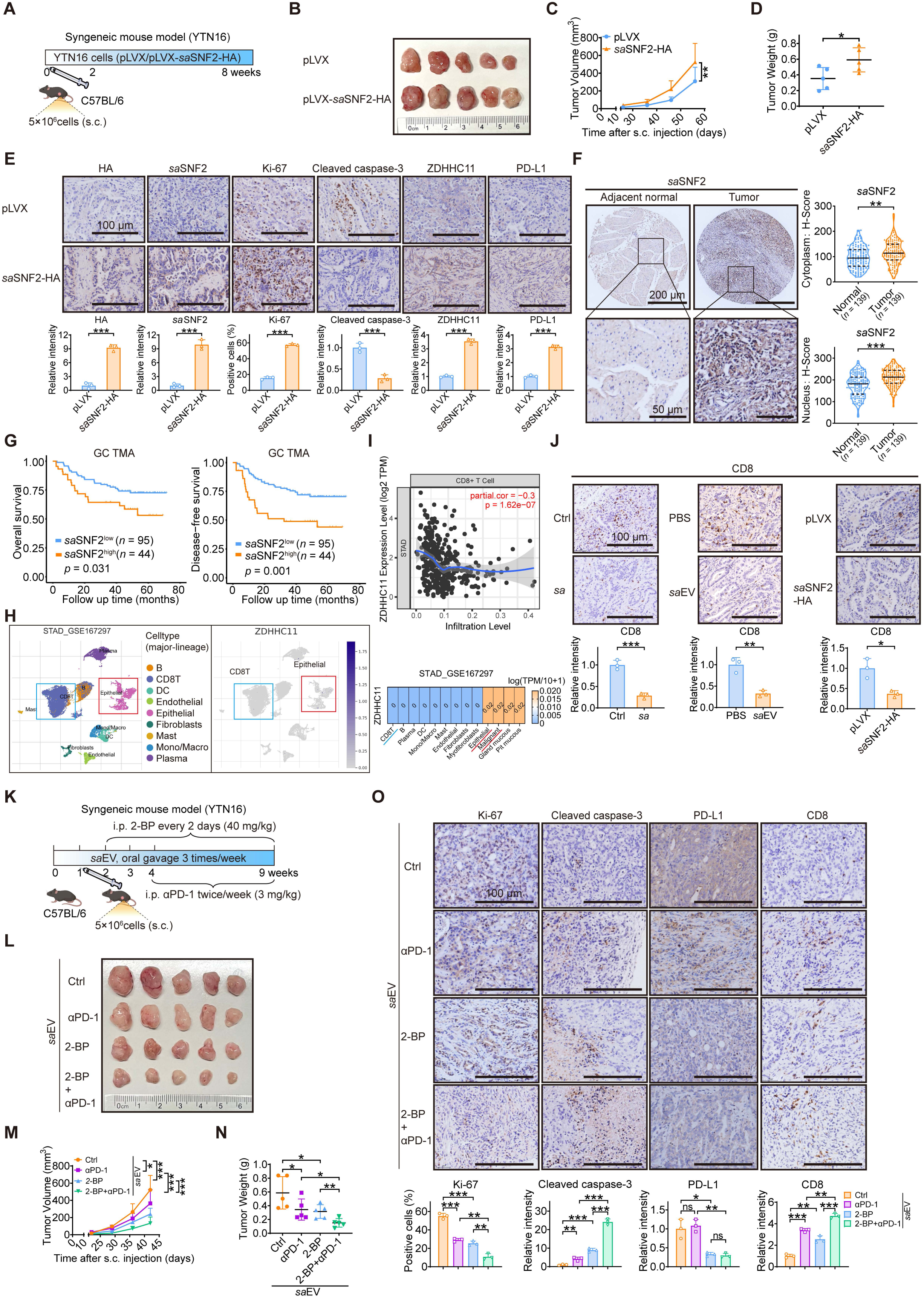
*sa*SNF2 is a bacterial tumor marker, and targeting the *sa*SNF2-ZDHHC11 pathway suppresses tumor growth and synergizes with anti-PD-1 therapy. (A) Schematic of the subcutaneous mouse model design. YTN16 cells with stable *sa*SNF2 overexpression were subcutaneously injected into C57BL/6 mice. (B) Illustrative images showing tumors from syngeneic mice (YTN16). (C and D) Data on tumor growth curves and harvested tumor weights are shown. (E) Tumor tissue sections were immunohistochemically stained for HA, *sa*SNF2, Ki-67, Cleaved caspase-3, ZDHHC11 and PD-L1 expression (upper panel). Staining intensity was quantified using ImageJ and presented as bar graphs (lower panel). (F) Clinical specimens were immunohistochemically stained for *sa*SNF2 expresstion (left panel). Staining intensity in the cytoplasm and nucleus was quantified separately based on the H-Score and is presented as a violin plot (*n* = 139) (right panel). (G) The prognostic significance of *sa*SNF2 for overall survival and disease-free survival was evaluated using Kaplan-Meier analysis (*n* = 139). (H) ZDHHC11 expression across different cell types in gastric cancer (GC) tissues, as analyzed by single-cell RNA sequencing (scRNA-seq) from the TISCH database (http://tisch.comp-genomics.org/). (I) Correlation between ZDHHC11 expression and CD8^+^ T cell infiltration in GC, analyzed using scRNA-seq data from the TIMER2.0 database (http://timer.cistrome.org/). (J) Tumor tissue sections were immunohistochemically stained for CD8 expression (upper panel). Staining intensity was quantified using ImageJ and presented as bar graphs (lower panel). (K) Schematic of the subcutaneous mouse model design. C57BL/6 mice received oral gavage of *sa*EVs three times weekly. Anti-PD1 (3 mg/kg, twice weekly) and 2-BP (40 mg/kg, every other day) were intraperitoneal (i.p.) injected into the mice. (L) Illustrative images showing tumors from syngeneic mice (YTN16). (M and N) Data on tumor growth curves and harvested tumor weights are shown. (O) Tumor tissue sections were immunohistochemically stained for Ki-67, Cleaved caspase-3, PD-L1 and CD8 expression (upper panel). Staining intensity was quantified using ImageJ and presented as bar graphs (lower panel). Data are presented as mean ± SD. \**p* < 0.05; \*\**p* < 0.01; \*\*\**p* < 0.001; ns, not significant. S.c., subcutaneous; TMA, tissue microarray; STAD, Stomach adenocarcinoma; EV, extracellular vesicle; 2-BP, 2-Bromohexadecanoic acid; i.p., intraperitoneal injection; αPD-1, anti-PD1.

To assess the clinical relevance of *sa*SNF2, we examined its expression in gastric cancer (GC) tissues and matched adjacent normal tissues. After generating and validating a monoclonal antibody against *sa*SNF2 (Figures S6A–S6E), we used it in conjunction with an HA-tag antibody to confirm *sa*SNF2-HA expression in syngeneic mouse tumors (Figure 7E). IHC analysis of 139 clinical specimens demonstrated significantly higher *sa*SNF2 expression in GC tissues relative to adjacent normal controls (Figure 7F). Kaplan–Meier survival analysis further revealed that elevated *sa*SNF2 expression correlated with worse overall and disease-free survival (Figure 7G) and was significantly associated with advanced TNM stage (Table S1). Together, these findings underscore the clinical importance of *sa*SNF2 in promoting gastric cancer progression.

### *SA*-mediated PD-L1 stabilization via palmitoylation in gastric cancer represents a potential therapeutic target, which may be exploited by combining palmitoylation inhibitors with anti-PD1 therapy

Given that the *sa*SNF2–ZDHHC11 axis regulates PD-L1 palmitoylation and stabilization, we hypothesized it may also modulate the tumor immune microenvironment. In a gastric cancer (GC) scRNA-seq dataset from TISCH (http://tisch.comp-genomics.org/), *ZDHHC11* expression was higher in epithelial and malignant cells than in tumor-infiltrating lymphocytes (TILs), particularly CD8^+^ T cells (Figure 7H). Analysis of an independent dataset from TIMER2.0 (http://timer.cistrome.org/) revealed a significant negative correlation between *ZDHHC11* expression and CD8^+^ T cell infiltration levels (Figure 7I), suggesting that ZDHHC11 may help establish an immunosuppressive niche. Consistently, in our experimental models, administration of *S. anginosus*, *sa*EVs, or *sa*SNF2 significantly reduced CD8^+^ T cell infiltration compared to respective controls (Figure 7J). Together, these results indicate that the *sa*SNF2–ZDHHC11 axis promotes an immunosuppressive tumor microenvironment by stabilizing PD-L1 and, consequently, attenuating CD8⁺ T cell infiltration.

Prompted by these findings, we investigated the effect of ZDHHC11 inhibition (using 2-BP) and/or anti-PD1 therapy in gastric cancer colonized with *SA*, given that *SA* upregulates ZDHHC11 to stabilize PD-L1. We established a mouse gastric cancer model by gavaging C57BL/6 mice with *sa*EV and evaluated the efficacy of 2-BP plus anti-PD1 on *SA*- promoted tumorigenesis (Figure 7K). While either 2-BP or anti-PD1 alone partially suppressed tumor growth in the presence of *sa*EV—as indicated by reduced tumor volume, weight, and Ki-67 expression (Figures 7L–7O)—the combination of 2-BP and anti-PD1 exerted a dramatically stronger anti-tumor effect (Figures 7L–7N). IHC staining confirmed that the combination effectively lowered PD-L1 expression and increased CD8⁺ T cell infiltration in the *sa*EV-induced model (Figure 7O).

These results suggest that targeting the *sa*SNF2–ZDHHC11–PD-L1 axis represents a rational therapeutic strategy for *SA*- enriched gastric cancer. Consequently, *SA*- regulated proteins such as ZDHHC11 and PD- L1 could serve as biomarkers to guide treatment, and inhibitors of ZDHHC11 activity and PD- L1 expression may represent novel therapeutic agents for *SA*-positive GC patients.

## Discussion

Our study reveals a previously unknown mechanism of cross-kingdom epigenetic regulation in gastric cancer (GC), driven by the oncobacterium *Streptococcus anginosus* (*SA*). We identify a bacterial chromatin remodeler, *sa*SNF2, which is delivered to host cancer cells via extracellular vesicles (*sa*EVs). Within the nucleus, *sa*SNF2 integrates into host chromatin-remodeling complexes to co-opt the transcriptional program, simultaneously driving tumor proliferation and establishing an immunosuppressive microenvironment (Figure 8). This work not only establishes *SA* as a pivotal GC driver but also uncovers the *sa*SNF2-ZDHHC11-PD-L1 axis as a novel therapeutic target to overcome immune evasion.

**Figure 8.**
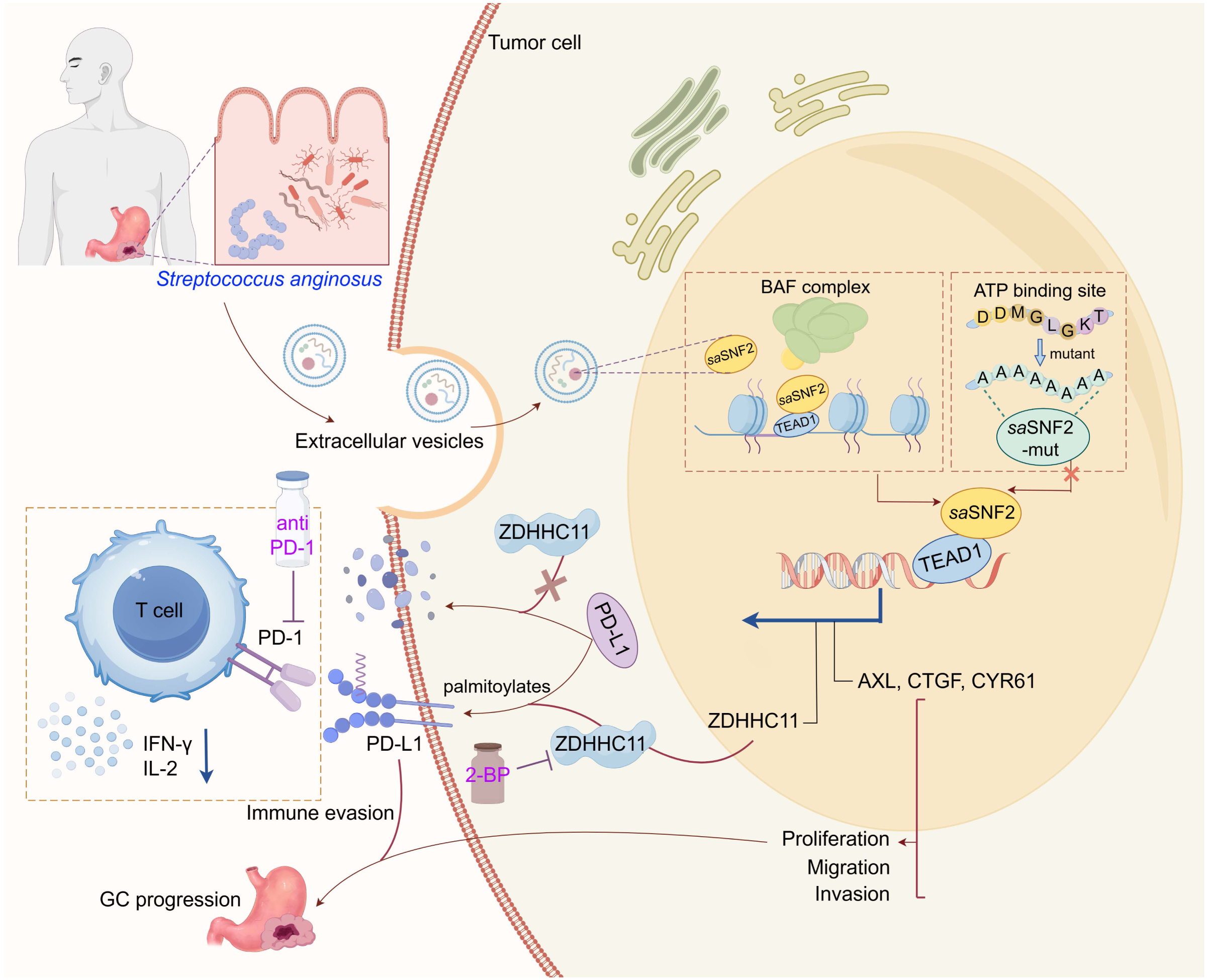
Scheme of *sa*SNF2/TEAD1/ZDHHC11/PD-L1 axis in promoting GC progression. Bacterial *sa*SNF2, delivered via *sa*EVs, interacts with host transcription factor TEAD1 to upregulate ZDHHC11 and other TEAD1 downstream target genes. ZDHHC11 palmitoylates and stabilizes PD-L1, driving immune evasion and tumor progression. Inhibition of ZDHHC11 restores anti-tumor immunity and enhances anti-PD-1 efficacy. The schematic was designed using Figdraw.

### *sa*EVs-activated ZDHHC11 is a modulator of PD-L1

Bacterial EVs play a critical role in communication with host cells by delivering proteins and nucleosides^26, 27^. Here, we show that *sa*EVs function as pivotal inter-kingdom shuttles, carrying a chromatin remodeler that induces expression of *ZDHHC11*. Given that the related enzymes ZDHHC3 and ZDHHC9 regulate PD-L1 palmitoylation and stabilization^15, 28^, we hypothesized that ZDHHC11—a member of the same protein acyltransferase family^29^—might similarly modify PD-L1. Palmitoylation, the attachment of palmitate to cysteine residues, critically regulates membrane protein interactions^30^. Our results confirm that ZDHHC11 does indeed palmitoylate PD-L1, uncovering a previously elusive mechanism of *sa*EV-mediated immune evasion in gastric cancer.

We established a structural model to predict how ZDHHC11 recognizes and binds PD-L1. This model identified key binding residues (Ser54, Ser98, and Arg165) within the enzyme’s ankyrin repeat domain (Figure 3F). This parallels the known function of ankyrin repeats in related enzymes like ZDHHC13 and ZDHHC17, which bind Huntington (HTT) protein for palmitoylation^31, 32^. Through this binding, ZDHHC11 palmitoylates PD-L1, stabilizing it to enhance surface expression, suppress T cell activity, and promote immune evasion. Thus, blocking PD-L1 palmitoylation mitigates its expression and restores T cell cytotoxicity.

### *sa*EV-derived *sa*SNF2 functions as a cross-kingdom transcriptional co-activator

We demonstrated that *sa*SNF2 enhances the activity of the transcriptional enhancer factor TEF-1 (TEAD1), which in turn acts as a transcriptional activator of *ZDHHC11*. This finding is particularly significant because TEAD1 is highly expressed in gastric cancer^33^. Furthermore, TEAD transcription factors are established prognostic biomarkers in breast, ovarian and prostate cancers^34^ and drive oncogenesis by activating downstream targets such as *AXL*, *CTGF*, *CYR61*, *SNAI1*, *MYC*, and *GLI2*^17,34^. Critically, we show that *sa*SNF2-activated TEAD1 directly mediates the transcription of *AXL*, *CTGF*, *CYR61*, and *SNAI1* to promote gastric cancer proliferation. These results provide important mechanistic support for the oncogenic role of *SA*.

We profiled the proteome of *sa*EVs to identify proteins capable of modulating host transcription and oncogenesis. Based on this analysis, we selected the chromatin remodeler *sa*SNF2 for further study. This choice was motivated by its homology to the human ATPase BRG1 (SMARCA4), a core subunit of the oncogenic SWI/SNF (BAF) chromatin remodeling complex. BRG1 utilizes ATP hydrolysis to slide or eject nucleosomes, thereby controlling DNA accessibility for transcription factors^35^ and is a poor prognostic marker in several cancers^36,38^.

This homology raised a key question: could *sa*SNF2 function cross-kingdom to similarly remodel chromatin and activate mammalian transcription? We found that *sa*SNF2 does indeed physically associate with the transcriptional enhancer TEAD1. This interaction facilitates TEAD1-mediated activation of the *ZDHHC11* gene. Crucially, this function depends on *sa*SNF2’s ATPase activity, as mutations in its catalytic site abolished the enhancement of *ZDHHC11* expression (Figure 6A). This mechanism mirrors that of its human homolog BRG1, which uses ATP hydrolysis within the BAF complex to create chromatin accessibility and drive gene expression^37,38^. Thus, *sa*SNF2 acts as a prokaryotic chromatin remodeler that co-opts the host’s transcriptional machinery to activate specific oncogenic pathways.

### *sa*SNF2 remodels host BAF complexes through its ATPase activity to amplify oncogenic transcription

Although approximately 1400 distinct mammalian SWI/SNF (BAF) complex assemblies exist in various tissues^25^, how the bacterial protein *sa*SNF2 influences their assembly and subunit stoichiometry in gastric cancer cells remains unknown. Our analyses reveal that *sa*SNF2 alters the assembly of at least three key BAF complex subunits compared to controls or an ATPase-deficient *sa*SNF2 mutant (Figure 6L). Specifically, the distribution/association of ARID1A, BAF60A, and BAF47 with the host various BAF complexes increased in the presence of functional *sa*SNF2, but not its mutant. In contrast, the distribution/association of host BRG1 with the host various BAF complexes remained unchanged, indicating that *sa*SNF2 does not displace the host ATPase subunit. Given that ATP hydrolysis by SWI/SNF ATPases drives DNA translocation along nucleosomes (∼1 bp per ATP) ^25^, we propose that *sa*SNF2 (using its own ATPase activity) may supplement host BRG1 activity, accelerating DNA remodeling. This enhanced activity likely alters BAF complex composition, thereby amplifying the output of specific transcription factors.

It is important to note that the C-terminal domain (CTD) of BAF47—a core subunit of the BAF complex—contains a basic α helix that binds directly to the nucleosomal acidic patch^24^. This anchors the complex to chromatin. Meanwhile, the BAF60 subunit functions as a critical adaptor that links specific transcription factors to the remodeling complex to regulate gene expression^23^. The *sa*SNF2-dependent enhancement of BAF47 and BAF60 distribution/association with the host various BAF complexes likely amplifies this circuitry, thereby potentiating the activity of specific transcription factors. Finally, we show that TEAD1 is constitutively associated with the BAF complex; its association is only modestly altered by *sa*SNF2. This suggests that *sa*SNF2 primarily integrates into the existing host BAF machinery to exert its chromatin remodeling function, rather than radically altering TEAD1 recruitment. In summary, by engaging with and modulating the assembly of host various BAF complexes, *sa*SNF2 acts as a critical epigenetic regulator during tumorigenesis, exerting pleiotropic effects on cell cycle progression, immune response, and oncogenic signaling.

### *sa*SNF2 is a bacterial tumor marker, and palmitoylation inhibition plus anti-PD1 treatment can be a rational cancer therapy for gastric cancer colonized with tumor-resident *SA*

While many gastric cancer patients do not respond to PD-1/PD-L1 immune checkpoint blockade, the underlying mechanisms of resistance are poorly understood. A deeper knowledge of the pathways controlling PD-L1 expression and stability is therefore critical to improve therapeutic efficacy. Our study establishes a key molecular link between immune surveillance and the TEAD1–ZDHHC11–PD-L1 axis. We identify the bacterial chromatin remodeler *sa*SNF2 as a pivotal factor that co-opts the host transcription factor TEAD1 to upregulate *ZDHHC11* in *SA*-infected gastric cancer. ZDHHC11 subsequently stabilizes PD-L1 via palmitoylation, revealing a novel immune evasion pathway.

This axis provides a strong mechanistic rationale for a combination therapy. While the ZDHHC11 inhibitor 2-BP or anti-PD1 monotherapy each shows efficacy, their combination results in superior tumor suppression and a marked increase in CD8^+^ T cell infiltration in mouse gastric tumor models. Thus, targeting this pathway has important therapeutic implications for managing gastric cancer colonized with tumor-resident *SA*.

Furthermore, our finding that *sa*SNF2 integrates into and modulates the host BAF chromatin-remodeling complex suggests additional therapeutic strategies. Possible inhibitors targeting *sa*SNF2-enhanced BAF assembly or activity, such as PFI-3 (a specific inhibitor targeting BRG1)^39^ or FHD-286, another inhibitor of BRG1 used in treating AML^37^, may also provide another therapeutic strategy for *SA*-colonized gastric cancer.

In summary, we delineate a complete cross-kingdom signaling axis from microbial colonization to immune evasion. The *SA*-derived chromatin remodeler *sa*SNF2 integrates into host BAF complexes, cooperates with TEAD1 to activate oncogenic and immunomodulatory transcription, and ultimately elevates ZDHHC11-mediated PD-L1 palmitoylation. This work expands our understanding of the tumor microbiome’s functional role from a passive association to an active epigenetic driver of disease. Targeting the *sa*SNF2-ZDHHC11 axis presents a novel precision medicine strategy to potentiate immunotherapy for a defined subset of gastric cancer patients.

## Materials and Methods

### Patient and tissue specimens

Gastric cancer (GC) specimens, along with matched adjacent non-tumorous tissues, were collected from the Department of Gastric Surgery at the Sixth Affiliated Hospital of Sun Yat-sen University. Additionally, a gastric cancer tissue microarray (TMA) was also used, comprising 139 cases with paired tumor and adjacent normal tissue samples, along with associated clinical and survival data.

### Cell culture and transfection

Human gastric cancer cell lines AGS cells were obtained from Meisen Chinese Tissue Culture Collections (MeisenCTCC), while MKN1 and MKN45 were acquired from Servicebio. These cells were cultured in RPMI 1640 medium (Gibco) supplemented with 10% (v/v) fetal bovine serum (FBS; Gibco). The murine gastric cancer cell line YTN16 was kindly provided by Prof. Sachiyo Nomura (University of Tokyo, Japan) and maintained under standard culture conditions as previously described^40^. HEK293T were obtained from the American Type Culture Collection (ATCC) and cultured in Dulbecco’s Modified Eagle’s Medium (DMEM; Gibco) containing 10% FBS. All cells were maintained at 37 °C in a humidified atmosphere containing 5% CO₂.

For transient transfection experiments, plasmids were introduced into the cells using Polyethylenimine MAX 40,000 (Polysciences Inc., no. 24765-1) following the manufacturer’s protocol.

### Bacterial strain and cell co-culture

The *Streptococcus anginosus* strain (ATCC 33397) was obtained from ATCC. Bacterial cultures were maintained on Brain Heart Infusion (BHI) agar plates and grown aerobically at 37°C overnight in BHI broth.

For co-culture experiments, tumor cells were seeded in 60 mm culture dishes and allowed to adhere until reaching 70–90% confluence. *SA* was collected during the logarithmic growth phase. The bacterium-to-cell ratio was determined using multiplicity of infection (MOI), with an MOI of 30 being routinely employed for in vitro studies.

### Purification of extracellular vesicles (EVs)

EVs were isolated from *SA* cultures using an optimized ultracentrifugation protocol as described previously^41^. In brief, bacterial cultures were grown aerobically in 180 mL brain heart infusion (BHI) at 37°C until reaching logarithmic phase. Following cultivation, bacterial cells were removed by sequential centrifugation at 10,000 × g for 20 min (4°C) and filtration through 0.22 μm pore-size membranes (Millipore). The clarified supernatant was ultracentrifuged at 170,000 × g for 3 h at 4°C using an ultracentrifuge (Beckman Coulter, USA). The resulting EV pellet was resuspended in 300 μL PBS for subsequent experiments.

### EV labeling and cellular uptake analysis

*Streptococcus anginosus*-derived EVs were fluorescently labeled by incubation with 3,3’-dioctadecyloxacarbocyanine perchlorate (DiO; 20 μg/mL; MCE) for 40 min at 37°C. Unbound dye was removed by diafiltration against PBS using 30 kDa Amicon Ultra centrifugal filters (Merck Millipore) at 4,000 × g for 45 min.

Gastric cancer cells cultured in 12-well chamber slides were incubated with DiO-labeled EVs or sham controls for specified durations to assess cellular uptake. After PBS washing, cells were fixed with 4% paraformaldehyde (20 min), counterstained with DAPI (15 min), and imaged by confocal microscopy.

### Reagents and inhibitors

Endocytosis inhibitors were obtained from Aladdin and applied at the following working concentrations: chlorpromazine (5 μM, a clathrin-mediated endocytosis inhibitor), cytochalasin D (2.5 μM, an actin-mediated endocytosis inhibitor), and dynasore (40 μM, a dynamin-mediated endocytosis inhibitor). Protein palmitoylation modulators, including 2-bromopalmitate (2-BP; 50 μM for in vitro studies; 40 mg/kg for in vivo administration) and palmostatin B (5 μM), were acquired from Sigma-Aldrich.

### Syngeneic mouse model

Animal studies were conducted in specific-pathogen-free (SPF) laminar-flow ventilated cages under a 12/12-hour light/dark cycle. The syngeneic mouse model was established using 6-week-old male C57BL/6 mice. YTN16 cells (5 × 10^6^ cells/tumor) suspended in 50% Matrigel (Beyotime) were subcutaneously implanted into the mice. For bacteria administration, mice were orally gavaged with *SA* (1 × 10^8^ CFU suspended in 100 μl PBS) three times per week. For extracellular vesicles (EVs) treatment, mice were orally administered EVs (100 ug/mouse in 100 μl PBS) three times per week throughout the experimental period. For anti-PD1 treatment, mice received intraperitoneal injections of anti-PD1 antibody (3 mg/kg; BioXcell) twice weekly. For 2-BP treatment, mice received intraperitoneal injections of 2-BP (40 mg/kg; Sigma-Aldrich) every two days. Tumor dimensions were monitored regularly, with volume calculated using the formula: Volume (mm^3^) = (length × width^2^)/2. At the experimental endpoint, all tumors were surgically resected and weighed.

### 16s rRNA sequencing and data analysis

Fecal samples were collected from mice in both the control and *SA* gavage groups and subjected to 16S rRNA sequencing at seven distinct time points. The V3-V4 hypervariable region of bacterial 16S rRNA genes was amplified using primer pair 338F (5’-ACTCCTACGGGAGGCAGCA) and 806R (5’-GGACTACHVGGGTWTCTAAT), with unique 7-base barcodes added to enable sample multiplexing. Each PCR reaction mixture included: 5 μl of 5× reaction buffer, 0.25 μl Fast pfu DNA Polymerase (5 U/μl), 2 μl dNTPs (2.5 mM), 1 μl each of forward and reverse primers (10 μM), 1 μl DNA template, and 14.75 μl sterile distilled water. The thermal profile involved: 5 min initial denaturation at 98°C; 25 cycles of 98°C for 30 s, 53°C for 30 s, and 72°C for 45 s; followed by 5 min final extension at 72°C. PCR products were purified using Vazyme VAHTSTM DNA Clean Beads (Nanjing, China) and quantified with the Quant-iT PicoGreen dsDNA Assay Kit (Invitrogen, USA). Equimolar concentrations of purified amplicons were combined and subjected to paired-end sequencing (2250 bp) on an Illumina NovaSeq platform using the NovaSeq 6000 SP Reagent Kit (500 cycles) at Shanghai Personal Biotechnology Co., Ltd.

### Fluorescence in situ hybridization (FISH)

Bacterial colonization in human GC tissues was assessed by FISH using an FITC-labeled *Streptococcus anginosus*-specific probe (5’-/FITC/TCAAGCATCTAACATGTGTTACATACTG-3’). The FISH assay was performed according to the manufacturer’s protocol (EXONBIO, D-0016). Tissue localization of bacteria was visualized using a confocal microscopy.

### Cell proliferation and colony formation assays

For cell proliferation analysis, 15,000 cells per well were plated in 12-well plates. Continuous monitoring was performed at 4-hour intervals using a live-cell imaging system (Incucyte S3, Sartorius, Germany). The acquired images were processed, and data analysis was conducted using the manufacturer’s software (Incucyte 2021C). Quantitative data were extracted using the integrated analysis software (Incucyte 2019B Rev2).

In parallel, colony formation was assessed by seeding 1000 cells per well in 6-well plates, followed by incubation in culture medium for 7–14 days. At the end of the experiments, the colonies were fixed (4% paraformaldehyde, 15 min), stained (0.5% crystal violet, 30 min), and quantified.

### Cell migration and invasion assays

Cell migration and invasion assays were performed using Transwell chambers (Corning, NY, 353097) equipped with 8-μm pore size membranes as previously described^42^. For invasion assay, membranes were pre-coated with 100 µL of Matrix-Gel™ Basement Membrane Matrix (1:10 dilution, Beyotime, C0372) and incubated at 37 °C for 2 hours. Cells were suspended in serum-free medium at densities of 1 × 10⁵ (migration) or 1.5 × 10⁵ (invasion) and seeded into the upper chamber, while the lower chamber contained medium supplemented with 10% FBS. Cells were incubated for approximately 22 hours, then fixed (4% paraformaldehyde, 15 min) and stained (0.5% crystal violet, 30 min). Non-migrated/invaded cells were removed by swabbing. Cell counts were performed under a light microscope across ≥3 random fields per membrane, with triplicate replicates per experiment.

### Immunofluorescence (IF) staining

For tissue immunofluorescence staining, frozen tissue sections were equilibrated to room temperature for 15 min, fixed with 4% paraformaldehyde (PFA) for 30 min, blocked with 5% BSA in PBS for 1 h, and then incubated with primary antibodies overnight at 4°C. Alexa Fluor-conjugated secondary antibodies (Invitrogen) were applied for 1 h (RT, dark), followed by DAPI counterstaining (10 min). Slides were imaged using a confocal microscope.

For cell immunofluorescence staining, cells grown on 12-well chamber slides were fixed in 4% PFA (15 min), permeabilized with 0.5% Triton X-100 in 5% BSA/PBS (1 h, RT), and incubated with primary antibodies overnight at 4°C. After washing, secondary antibodies (Alexa Fluor-conjugated, Invitrogen) were applied (1 h, RT, dark), followed by DAPI staining. Images were acquired using a confocal microscope.

### RNA extraction and quantitative real-time PCR

Total RNA was isolated from cells with Total RNA Extractor (Servicebio) and reverse-transcribed into cDNA using GoScript™ Reverse Transcription Kit (Promega) according to the manufacturer’s protocol. Quantitative real-time PCR was conducted on a LightCycler® 480 II system (Roche) with SYBR Green qPCR Master Mix (TransGen Biotech). All reactions were performed in triplicate using Actin as the endogenous control.

### Immunoblot analyses and immunoprecipitation

Cells were harvested and lysed in lysis buffer (50 mM Tris-HCl [pH 7.5], 150 mM NaCl, 0.1% Triton X-100, 0.1% NP-40, 1 mM EDTA) supplemented with protease and phosphatase inhibitors (Bimake). Protein concentrations were determined by BCA assay (Servicebio), with equal amounts separated by SDS-PAGE and transferred to PVDF membranes (Millipore). After overnight incubation with primary antibodies at 4°C, membranes were incubated with HRP-conjugated secondary antibodies (ThermoFisher; 1:10,000) for 1 h at room temperature. Protein bands were detected by enhanced chemiluminescence (ECL) and visualized on X-ray films.

For in vivo exogenous immunoprecipitation, cells pretreated with 25 μM MG132 (MedChemExpress) for 6 h were lysed in ice-cold lysis buffer. Cell lysates were immunoprecipitated using anti-Flag M2 agarose beads (Sigma-Aldrich) at 4°C overnight. Beads were washed four times with lysis buffer and eluted by boiling in 2× SDS loading buffer (10 min). Immunoblotting was performed with specified antibodies.

### Mass spectrometry and data processing

MKN1 cells transiently transfected with Flag-tagged PD-L1 plasmids were treated with *sa*EVs or PBS (control). PD-L1 was immunoprecipitated using anti-Flag M2 beads. After washing, bound proteins were eluted and separated by SDS-PAGE. The protein bands corresponding to PD-L1 were excised from the gel and sent to Cosmos Wisdom (Hangzhou, China) for further processing. Briefly, the gel slices were destained (50% acetonitrile/50 mM NH4HCO3), and subjected to reduction (10 mM TCEP, 37°C, 30 min) and alkylation (25 mM iodoacetamide, RT, 30 min, dark). After washing, in-gel tryptic digestion (2 μg trypsin, 37°C, overnight) was performed, followed by peptide extraction (50% acetonitrile/0.1% formic acid) and vacuum drying. Peptides were separated via nanoLC (EASY-nLC 1200, Thermo Scientific) using a 60-min gradient (2–100% mobile phase B: 0.1% formic acid/80% acetonitrile, 400 nL/min) and analyzed by Q Exactive HF-X mass spectrometer (Orbitrap resolution: 60,000 [MS], 30,000 [MS/MS]; NCE: 27; data-dependent top20 mode). Data were processed using Proteome Discoverer 2.4 (human SwissProt 2023_09 database) with fixed (carbamidomethylation) and variable (oxidation, N-terminal acetylation, palmitoylation) modifications (precursor tolerance: 10 ppm; fragment tolerance: 0.01 Da).

### Acyl-biotin exchange (ABE) palmitoylation assay

AGS cells were transiently transfected with Flag-tagged PD-L1 constructs for 48 h. Following transfection, cell lysates were subjected to immunoprecipitation with anti-Flag M2 agarose beads overnight at 4°C. Protein palmitoylation was then assessed using an acyl-biotinyl exchange (ABE) assay (Immuno-Precipitation-ABE Palmitoylation Kit for WB, AM10314; AIMSMASS Co., Ltd., Shanghai, China) following the manufacturer’s protocol.

### Turnover assay

Cells were seeded in six-well plates and treated with cycloheximide (CHX; Sigma) at a final concentration of 100 μg/mL. At specified time points post-treatment, cells were harvested for immunoblot analysis. Protein levels were quantified using ImageJ software.

### Ubiquitination assay

Gastric cancer cells were co-transfected with a His-tagged ubiquitin plasmid along with the specified expression constructs. Following a 48-h incubation, cells were treated with the proteasome inhibitor MG132 (25 μM; MedChemExpress) for 6 h prior to collection. Subsequently, cells were harvested and lysed in denaturing buffer A (6 M guanidine-HCl, 0.1 M Na2HPO4/NaH2PO4, 10 mM imidazole, pH 8.0) followed by brief sonication. The lysates were then incubated with Ni-NTA agarose beads (Invitrogen) at 4 °C overnight with gentle rotation. The beads were sequentially washed as follows: (1) buffer A (×3), (2) a 1:3 mixture of buffer A and buffer TI (25 mM Tris-HCl, 20 mM imidazole, pH 6.8) (×2), and (3) buffer TI (×2). The eluted proteins were subjected to SDS-PAGE and analyzed via immunoblotting.

### In situ Proximity Ligation Assay (PLA)

The PLA assay was conducted using the Duolink® PLA kit (Sigma-Aldrich, DUO92101) following the manufacturer’s protocol. Briefly, 293T cells were co-transfected with HA-tagged SNF2 and Flag-tagged TEAD1 expression plasmids. After 24 hours, cells were fixed with 4% paraformaldehyde for 10 min at room temperature, followed by permeabilization with 0.5% Triton X-100 for 10 min. Subsequently, cells were blocked with the provided blocking buffer for 1 hour at room temperature, followed by incubation with primary antibodies against HA (rabbit polyclonal, Proteintech, 51064-2-AP) and Flag (mouse monoclonal, Sigma, F1804) at 4°C overnight. Thereafter, cells were treated with PLA secondary probe for 1 hour at 37°C. The ligation reaction was performed by applying the ligation mixture (37°C, 30 min), followed by amplification using the polymerization mix (37°C, 100 min). Post-washing, samples were mounted with Duolink® In Situ Mounting Medium containing DAPI and imaged using a confocal microscope.

### Immunohistochemical (IHC) staining

The immunohistochemical staining was performed as previously described^43^. Tissue sections embedded in paraffin underwent deparaffinization using xylene and gradual ethanol hydration. Antigen retrieval was performed by microwave heating in sodium citrate buffer (pH 6.0). After cooling, endogenous peroxidase activity was quenched with 3% H2O2 (10 min, RT), followed by 1-hour blocking with goat serum (RT). Primary antibody incubation proceeded overnight at 4°C, with subsequent 20-minute RT incubation using secondary antibodies. Following diaminobenzidine (Zhongshan Goldenbridge Biotechnology) visualization, sections were hematoxylin-counterstained, dehydrated, and coverslipped. Staining scores were determined by calculating either the percentage score or the intensity score.

### OVA-specific CD8^+^ T cell isolation and co-culture experiments

CD8^+^ T cells were isolated from spleens of OT-1 TCR transgenic mice (C57BL/6J background) using a MojoSort™ Mouse CD8 T Cell Isolation Kit (BioLegend, 480007). The isolated cells were then cultured in RPMI 1640 medium supplemented with 10% fetal bovine serum, 1% penicillin/streptomycin, 2 mM L-glutamine, 50 μM 2-mercaptoethanol, 10 ng/ml recombinant mouse IL-2 (BioLegend, 575402), and 100 nM OVA257-264 peptide (MCE, HY-P1489) for primary stimulation. After 48 hours of activation and expansion, cells were washed and maintained in peptide-free medium for an additional 48-hour resting period.

For functional assays, activated CD8^+^ T cells were co-cultured with YTN16-OVA cells for 24 hours. Culture supernatants were collected for quantification of IL-2 and IFN-γ secretion using commercial ELISA kits (Servicbio), while CD8^+^ T cells were harvested simultaneously for RNA extraction and subsequent qPCR analysis of *Il2* and *Ifng* transcript levels. YTN16-OVA cell viability was monitored by measuring the cell index, which was normalized to baseline values at co-culture initiation (time = 0 h). As a control, YTN16-OVA cells cultured alone under identical conditions were included to assess cell death.

### Chromatin immunoprecipitation (ChIP)

The ChIP assay was performed as described^44^. Briefly, cells were cultured in 150-mm dishes, and then cross-linked with 1% formaldehyde for 20 min at room temperature. The reaction was quenched using 0.125 M glycine. After lysis in ChIP lysis buffer, the samples were centrifuged, and the pellet was resuspended in nuclear lysis buffer before sonication (Diagenode Bioruptor Pico sonicator; 30 sec on/off cycles, 20 cycles total). A 10-μl aliquot of sheared chromatin was reserved as input control, while the remainder was incubated overnight at 4 °C with either anti-Flag antibody (Sigma) or control rabbit IgG, coupled to ChIP beads. The immunoprecipitated chromatin underwent sequential washes with low-salt, high-salt, LiCl, and Tris-EDTA buffers. After protein-DNA cross-links were reversed by proteinase K treatment, the DNA was isolated with a PCR purification kit (Omega). Enrichment at TEAD1-binding sites was quantified by qRT-PCR.

### Protein Structure Preparation and Molecular Docking

The target protein sequence and structure were retrieved from the UniProt database (https://www.uniprot.org/). The obtained structure was imported into Discovery Studio 2019 for structural optimization. The optimized protein structure was then used for molecular docking. Protein-protein docking was performed using GRAMM (https://gramm.compbio.ku.edu/). The docking results were visualized and analyzed using PyMOL 3.1.

### Molecular Dynamics (MD) simulations

Molecular dynamics (MD) simulations were conducted using GROMACS 2022 software. The protein-ligand systems were prepared with the CHARMM36^45^ force field for proteins and GAFF2 for ligands, solvated in a TIP3P water box (1.2 nm periodic boundary). Long-range electrostatics were treated using the Particle Mesh Ewald (PME) method, while the Verlet algorithm handled short-range interactions. The system was first equilibrated through 100,000 steps of simulation in both the NVT (isothermal-isochoric) and NPT (isothermal-isobaric) ensembles, using a coupling time constant of 0.1 ps over a 100 ps period. A cutoff distance of 1.0 nm was applied for computing van der Waals and Coulomb interactions. Subsequently, a production MD run was performed using Gromacs 2022 under constant temperature (310 K) and pressure (1 bar) conditions for a total simulation time of 100 ns.

### Gel filtration chromatography

AGS cells were transiently transfected with either HA-tagged wild-type *sa*SNF2 (HA-*sa*SNF2-WT) or its mutant variant (HA-*sa*SNF2-mut) expression plasmids. 48 hours post-transfection, cells were harvested and lysed in lysis buffer containing 50 mM Tris–HCl (pH 7.5), 150 mM NaCl, 0.1% NP-40, 0.1% Triton X-100, and 1 mM EDTA, supplemented with protease and phosphatase inhibitor cocktail. The lysates were loaded onto a size-exclusion Superose 6 column (GE). Chromatography was performed using the GE AKTA avant150 chromatography system at 4°C with a flow rate of 0.4 mL/min. Fractions (400 μL each) were collected and subsequently analyzed by SDS-PAGE, followed by immunoblotting with the indicated antibodies.

### Monoclonal antibody (anti-*sa*SNF2) production via hybridoma technology

The *sa*SNF2-specific antibody was developed by Sino Biological Inc. (Beijing, China). To generate the antibody, Balb/c mice were immunized with VLP-conjugated peptides. Following four protein and three peptide immunizations, serum ELISA confirmed immunogenicity. Splenocytes from high-titer mice were then fused with SP2/0 myeloma cells, and hybridoma screening yielded antigen-specific clones. These clones were expanded and purified via Protein A affinity chromatography, achieving >90% purity. Binding specificity was validated by ELISA.

### Statistical analysis

All experiments were repeated a minimum of three times, with data shown as mean ± SD. Statistical comparisons between two groups were conducted using a two-tailed Student’s *t*-test, while multiple groups were evaluated by one-way ANOVA. For paired samples, a paired *t*-test was employed, and growth curves were analyzed via two-way ANOVA. Kaplan–Meier survival curves were assessed using the log-rank test. A value of *p* < 0.05 was set for statistical significance (\**p* < 0.05, ** *p* < 0.01, *** *p* < 0.001). Further details on statistical methods are provided in the respective figure legends. Data analysis was performed using SPSS and GraphPad Prism software.

## Acknowledgments

This work was supported by the National Key R&D Program of China (2020YFA0803300), the National Natural Science Foundation of China (82573782, 82273133, 82373139 and 82303028), Guangdong Basic and Applied Basic Research Foundation (2023A1515030261, 2024A1515013206), the Guangzhou Science and Technology Program Project (202206010167), and the National Key Clinical Discipline, the program of Guangdong Provincial Clinical Research Center for Digestive Diseases (2020B1111170004).

## Author Contributions

Mong-Hong Lee and Xiangqi Meng conceived and designed the research. Shi Chen ascertained and processed clinical specimens. Xiaoshan Xie, Yue Wei, Zhikai Zheng performed most of the biochemical and molecular experiments, with the assistance from Jiaying Zheng, Xijie Chen, Jiarui Wang, Ning Ma, Xiaoling Huang, Peng Zhang, Boyu Zhang, Hanyong Cai and Li Ma. Xiaoshan Xie, Yue Wei, Zhikai Zheng and Xijie Chen performed mice experiments. Xiaoshan Xie and Zhikai Zheng conducted the bioinformatics analyses. Yue Wei, Zhikai Zheng and Qingxin Liu contributed to discussion and data interpretation. Lishi Xiao collected and processed clinical specimens. Wenyu Wang provided OT-1 mice. Sachiyo Nomura provided YTN16 cells. Xiaoshan Xie, Xiangqi Meng and Mong-Hong Lee wrote the manuscript. All authors have read and approved the final manuscript.

## Data Availability

The data reported are available in the article itself or in its online Supplementary Materials.

## Conflict of Interest

The authors declare no conflict of interest.

## Notes

### Competing Interest Statement

The authors have declared no competing interest.

## References

1. Bray F, et al. Global cancer statistics 2022: GLOBOCAN estimates of incidence and mortality worldwide for 36 cancers in 185 countries. CA: A Cancer Journal for Clinicians 74, 229–263 (2024).

2. Kumar S, Metz DC, Ellenberg S, Kaplan DE, Goldberg DS. Risk Factors and Incidence of Gastric Cancer After Detection of Helicobacter pylori Infection: A Large Cohort Study. Gastroenterology 158, 527–536.e527 (2020).

3. Zhang T, et al. Intratumoral Fusobacterium nucleatum Recruits Tumor-Associated Neutrophils to Promote Gastric Cancer Progression and Immune Evasion. Cancer Res 85, 1819–1841 (2025).

4. Fu K, et al. Streptococcus anginosus promotes gastric inflammation, atrophy, and tumorigenesis in mice. Cell 187, 882–896.e817 (2024).

5. Wang H, et al. Streptococcus lutetiensis inhibits CD8+ IL17A+ TRM cells and leads to gastric cancer progression and poor prognosis. npj Precision Oncology 9, (2025).

6. Lochhead P, El-Omar EM. Molecular Predictors of Gastric Neoplastic Progression. Cancer Cell 33, 9–11 (2018).

7. Guo F, Li L, Li L. Streptococcus anginosus: A new pathogen of superficial gastritis, atrophic gastritis and gastric cancer. Biomolecules & biomedicine 24, 1040–1043 (2024).

8. Yasuda T, Wang YA. Gastric cancer immunosuppressive microenvironment heterogeneity: implications for therapy development. Trends in cancer 10, 627–642 (2024).

9. Janjigian YY, et al. First-line nivolumab plus chemotherapy versus chemotherapy alone for advanced gastric, gastro-oesophageal junction, and oesophageal adenocarcinoma (CheckMate 649): a randomised, open-label, phase 3 trial. Lancet 398, 27–40 (2021).

10. Joshi SS, Badgwell BD. Current treatment and recent progress in gastric cancer. CA Cancer J Clin 71, 264–279 (2021).

11. Chong X, et al. Recent developments in immunotherapy for gastrointestinal tract cancers. J Hematol Oncol 17, 65 (2024).

12. Kang E-J, et al. The secreted protein Amuc_1409 from Akkermansia muciniphila improves gut health through intestinal stem cell regulation. Nature Communications 15, (2024).

13. Yaghoubfar R, et al. Effect of Akkermansia muciniphila, Faecalibacterium prausnitzii, and Their Extracellular Vesicles on the Serotonin System in Intestinal Epithelial Cells. Probiotics and antimicrobial proteins 13, 1546–1556 (2021).

14. Yao H, et al. Inhibiting PD-L1 palmitoylation enhances T-cell immune responses against tumours. Nature Biomedical Engineering 3, 306–317 (2019).

15. Li Z, Jiang D, Liu F, Li Y. Involvement of ZDHHC9 in lung adenocarcinoma: regulation of PD-L1 stability via palmitoylation. In Vitro Cellular & Developmental Biology - Animal 59, 193–203 (2023).

16. Zhou Y, Huang T, Cheng A, Yu J, Kang W, To K. The TEAD Family and Its Oncogenic Role in Promoting Tumorigenesis. International Journal of Molecular Sciences 17, (2016).

17. Kumar R, Hong W. Hippo Signaling at the Hallmarks of Cancer and Drug Resistance. Cells 13, (2024).

18. Dunne PD, et al. AXL is a key regulator of inherent and chemotherapy-induced invasion and predicts a poor clinical outcome in early-stage colon cancer. Clin Cancer Res 20, 164–175 (2014).

19. Chen YC, et al. Obesity-associated leptin promotes chemoresistance in colorectal cancer through YAP-dependent AXL upregulation. American journal of cancer research 11, 4220–4240 (2021).

20. Clapier CR, Iwasa J, Cairns BR, Peterson CL. Mechanisms of action and regulation of ATP-dependent chromatin-remodelling complexes. Nat Rev Mol Cell Biol 18, 407–422 (2017).

21. Park J, Kirkland JG. The role of the polybromo-associated BAF complex in development. Biochemistry and cell biology = Biochimie et biologie cellulaire 103, 1–8 (2025).

22. Xu C, et al. Comprehensive molecular phenotyping of ARID1A-deficient gastric cancer reveals pervasive epigenomic reprogramming and therapeutic opportunities. Gut 72, 1651–1663 (2023).

23. Wang RR, Pan R, Zhang W, Fu J, Lin JD, Meng ZX. The SWI/SNF chromatin-remodeling factors BAF60a, b, and c in nutrient signaling and metabolic control. Protein Cell 9, 207–215 (2018).

24. Valencia AM, et al. Recurrent SMARCB1 Mutations Reveal a Nucleosome Acidic Patch Interaction Site That Potentiates mSWI/SNF Complex Chromatin Remodeling. Cell 179, 1342–1356.e1323 (2019).

25. Gourisankar S, Krokhotin A, Wenderski W, Crabtree GR. Context-specific functions of chromatin remodellers in development and disease. Nat Rev Genet 25, 340–361 (2024).

26. Choi HI, et al. Helicobacter pylori-derived extracellular vesicles increased in the gastric juices of gastric adenocarcinoma patients and induced inflammation mainly via specific targeting of gastric epithelial cells. Experimental & molecular medicine 49, e330 (2017).

27. Ahmadi Badi S, Khatami SH, Irani SH, Siadat SD. Induction Effects of Bacteroides fragilis Derived Outer Membrane Vesicles on Toll Like Receptor 2, Toll Like Receptor 4 Genes Expression and Cytokines Concentration in Human Intestinal Epithelial Cells. Cell J 21, 57-61 (2019).

28. Wang Q, et al. Benzosceptrin C induces lysosomal degradation of PD-L1 and promotes antitumor immunity by targeting DHHC3. Cell reports Medicine 5, 101357 (2024).

29. Malgapo MIP, Linder ME. Substrate recruitment by zDHHC protein acyltransferases. Open biology 11, 210026 (2021).

30. Linder ME, Deschenes RJ. Palmitoylation: policing protein stability and traffic. Nat Rev Mol Cell Biol 8, 74–84 (2007).

31. Huang K, et al. Neuronal palmitoyl acyl transferases exhibit distinct substrate specificity. Faseb j 23, 2605–2615 (2009).

32. Lemonidis K, Sanchez-Perez MC, Chamberlain LH. Identification of a Novel Sequence Motif Recognized by the Ankyrin Repeat Domain of zDHHC17/13 S-Acyltransferases. J Biol Chem 290, 21939–21950 (2015).

33. Tanaka Y, et al. Multi-omic profiling of peritoneal metastases in gastric cancer identifies molecular subtypes and therapeutic vulnerabilities. Nat Cancer 2, 962–977 (2021).

34. Zhou Y, Huang T, Cheng AS, Yu J, Kang W, To KF. The TEAD Family and Its Oncogenic Role in Promoting Tumorigenesis. Int J Mol Sci 17, (2016).

35. Ribeiro-Silva C, Vermeulen W, Lans H. SWI/SNF: Complex complexes in genome stability and cancer. DNA repair 77, 87–95 (2019).

36. Huang LY, et al. SCF(FBW7)-mediated degradation of Brg1 suppresses gastric cancer metastasis. Nat Commun 9, 3569 (2018).

37. Fiskus W, et al. BRG1/BRM inhibitor targets AML stem cells and exerts superior preclinical efficacy combined with BET or menin inhibitor. Blood 143, 2059–2072 (2024).

38. Schick S, et al. Acute BAF perturbation causes immediate changes in chromatin accessibility. Nat Genet 53, 269–278 (2021).

39. Kang Q, et al. BRG1 promotes progression of B-cell acute lymphoblastic leukemia by disrupting PPP2R1A transcription. Cell Death Dis 15, 621 (2024).

40. Yamamoto M, et al. Established gastric cancer cell lines transplantable into C57BL/6 mice show fibroblast growth factor receptor 4 promotion of tumor growth. Cancer Science 109, 1480–1492 (2018).

41. Hong M, et al. Fusobacterium nucleatum aggravates rheumatoid arthritis through FadA-containing outer membrane vesicles. Cell Host Microbe 31, 798–810 e797 (2023).

42. Wei W, et al. FBXW7beta loss-of-function enhances FASN-mediated lipogenesis and promotes colorectal cancer growth. Signal Transduct Target Ther 8, 187 (2023).

43. Xie X, et al. EGF-Upregulated lncRNA ESSENCE Promotes Colorectal Cancer Growth through Stabilizing CAD and Ferroptosis Defense. *Research (Washington*, DC) 8, 0649 (2025).

44. Jin H, et al. The EIF3H-HAX1 axis increases RAF-MEK-ERK signaling activity to promote colorectal cancer progression. Nature Communications 15, (2024).

45. Jo S, Kim T, Iyer VG, Im W. CHARMM-GUI: a web-based graphical user interface for CHARMM. Journal of computational chemistry 29, 1859–1865 (2008).

