## Supplemental Figures and table for "A gastric microbial chromatin remodeler drives gastric cancer progression and immune evasion by reprogramming the host epigenome"

Figure S1

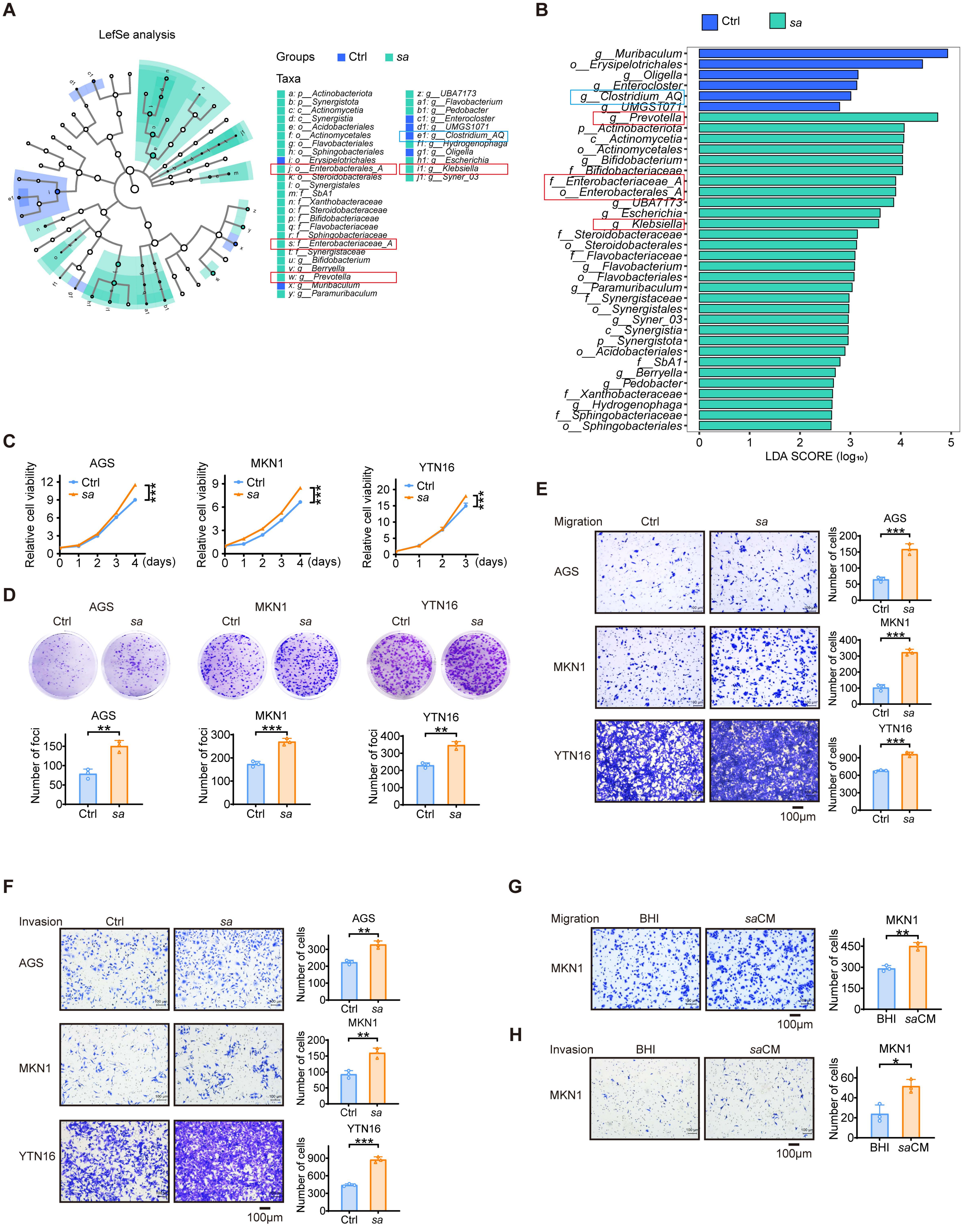

A
LefSe analysis
Groups Ctrl sa
Taxa
a: p__Actinobacteriota
b: p__Synergistota
c: c__Actinomycetia
d: c__Synergistia
e: o__Acidobacteriales
f: o__Actinomycetales
g: o__Flavobacteriales
h: o__Sphingobacteriales
i: o__Erysipelotrichales
j: o__Enterobacterales_A
k: o__Steroidobacterales
l: o__Synergistales
m: f__SbA1
n: f__Xanthobacteraceae
o: f__Steroidobacteraceae
p: f__Bifidobacteriaceae
q: f__Flavobacteriaceae
r: f__Sphingobacteriaceae
s: f__Enterobacteriaceae_A
t: f__Synergistaceae
u: g__Bifidobacterium
v: g__Berryella
w: g__Prevotella
x: g__Muribaculum
y: g__Paramuribaculum
z: g__UBA7173
a1: g__Flavobacterium
b1: g__Pedobacter
c1: g__Enterocloster
d1: g__UMGS1071
e1: g__Clostridium_AQ
f1: g__Hydrogenophaga
g1: g__Oligella
h1: g__Escherichia
i1: g__Klebsiella
j1: g__Syner_03
B
Ctrl
sa
g__Muribaculum
o__Erysipelotrichales
g__Oligella
g__Enterocloster
g__Clostridium_AQ
g__UMGS1071
g__Prevotella
p__Actinobacteriota
c__Actinomycetia
o__Actinomycetales
g__Bifidobacterium
f__Bifidobacteriaceae
f__Enterobacteriaceae_A
o__Enterobacterales_A
g__UBA7173
g__Escherichia
g__Klebsiella
f__Steroidobacteraceae
o__Steroidobacterales
f__Flavobacteriaceae
g__Flavobacterium
o__Flavobacteriales
g__Paramuribaculum
f__Synergistaceae
o__Synergistales
g__Syner_03
c__Synergistia
p__Synergistota
o__Acidobacteriales
f__SbA1
g__Berryella
g__Pedobacter
f__Xanthobacteraceae
g__Hydrogenophaga
f__Sphingobacteriaceae
o__Sphingobacteriales
0 1 2 3 4 5
LDA SCORE (log10)
C
AGS
MKN1
YTN16
Relative cell viability
Ctrl
sa
***
(days)
D
AGS
MKN1
YTN16
Ctrl
sa
Number of foci
**
***
**
E
Migration
Ctrl
sa
AGS
MKN1
YTN16
Number of cells
***
100μm
F
Invasion
Ctrl
sa
AGS
MKN1
YTN16
Number of cells
**
***
100μm
G
Migration
BHI
saCM
MKN1
Number of cells
**
100μm
H
Invasion
BHI
saCM
MKN1
Number of cells
*
100μm

**Figure S1. *SA* is a tumor-resident oncobacterium that drives gastric carcinogenesis.**

(A) LEfSe analysis (LDA score [$\log_{10}$] > 2) of microbial comparison in mice fecal samples following oral gavage with *SA*.

(B) Microbiota composition analyzed by linear discriminant analysis (LDA score [$\log_{10}$] > 2).

(C and D) The proliferative capacity and colony formation of AGS, MKN1 and YTN16 cells were examined after coculture with *SA*.

(E and F) The migratory and invasive capacities of AGS, MKN1 and YTN16 cells were assessed by Transwell assay following coculture with *SA*.

(G and H) The migratory and invasive capacities of MKN1 cells were assessed by Transwell assay after treatment with *SA*-conditioned medium (*sa*CM). Control was treated with BHI.

Data are presented as mean ± SD. $*p < 0.05$; $**p < 0.01$; $***p < 0.001$; ns, not significant. Ctrl, control; BHI, Brain heart Infusion Broth; CM, conditioned medium.

Data are presented as mean ± SD. $*p < 0.05$; $**p < 0.01$; $***p < 0.001$; ns, not significant. Ctrl, control; BHI, Brain heart Infusion Broth; CM, conditioned medium.

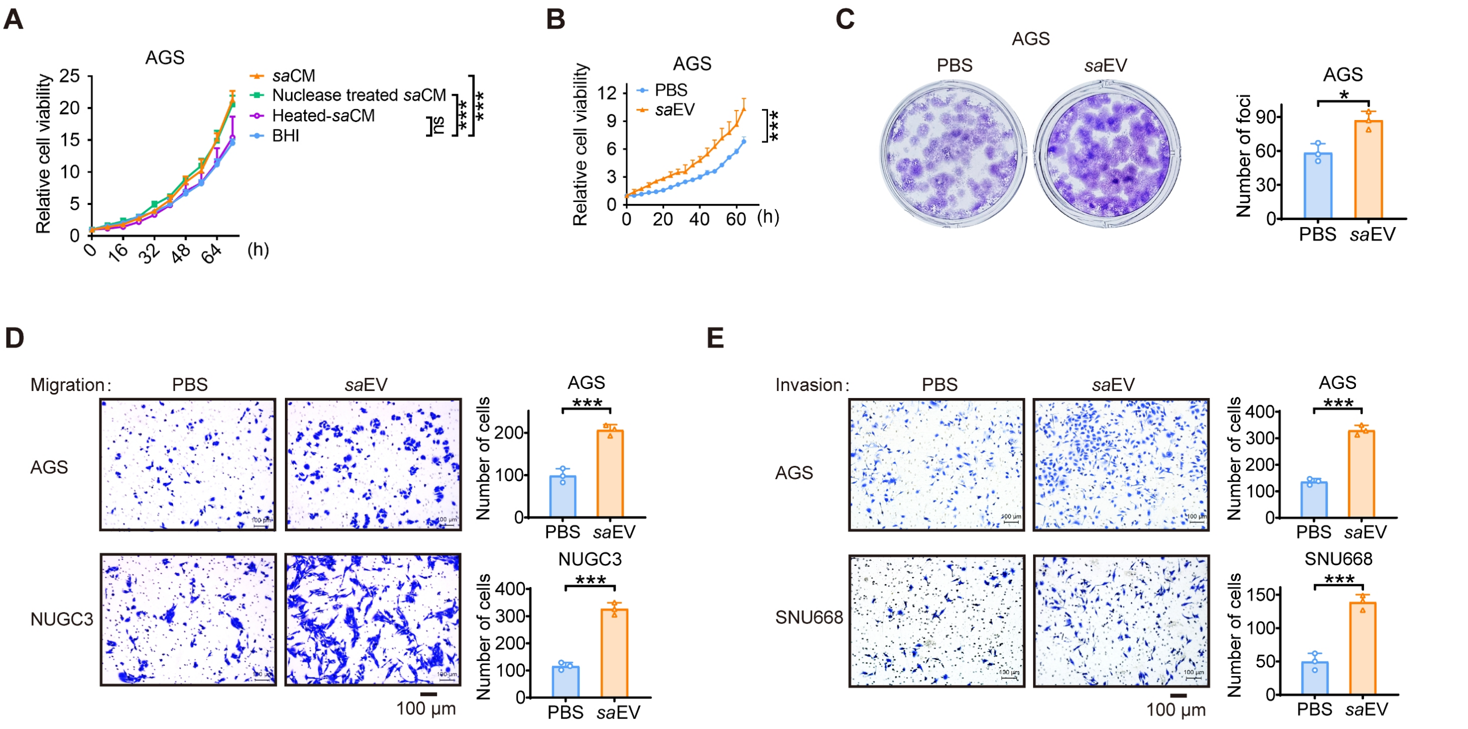

**Figure S2.** ***SA*** **extracellular vesicles drive tumor progression via dynamin-dependent endocytosis.**

(A) The proliferative capacity of AGS cells were examined after treatment with *sa*CM, nuclease treated *sa*CM or heated *sa*CM. Control was treated with BHI.

(B and C) The proliferative capacity and colony formation of AGS cells were examined after treatment with *sa*EVs. Control was treated with PBS.

(D and E) The migratory and invasive capacities of GC cells were assessed by Transwell assay after treatment with *sa*EVs. Control was treated with PBS.

Data are presented as mean ± SD. $*p < 0.05$; $**p < 0.01$; $***p < 0.001$; ns, not significant. BHI, Brain Heart Infusion Broth; CM, conditioned medium; EV, extracellular vesicle.

Figure S3

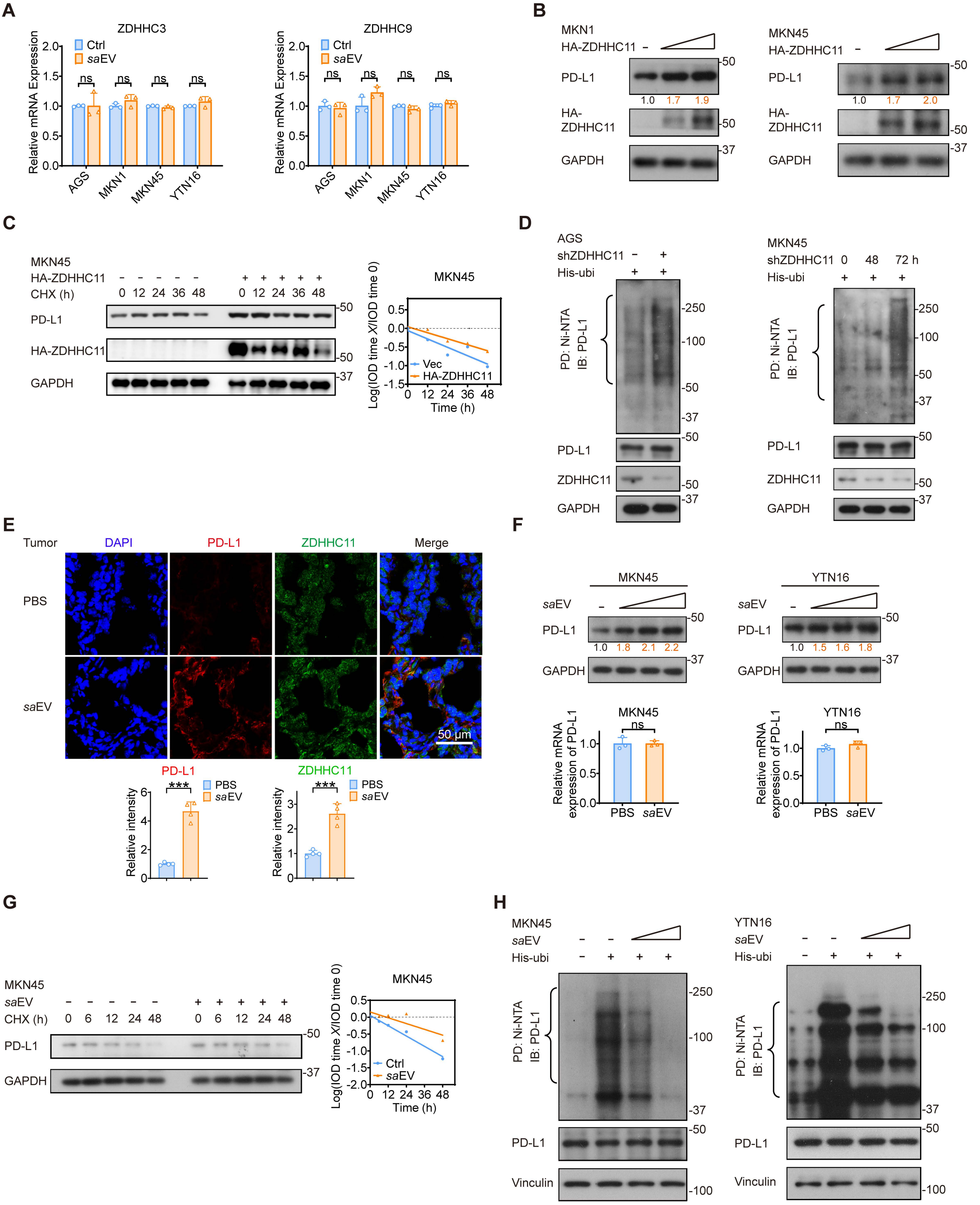
A
ZDHHC3
ZDHHC9
Ctrl
saEV
ns
Relative mRNA Expression
AGS
MKN1
MKN45
YTN16
B
MKN1
MKN45
HA-ZDHHC11
PD-L1
HA-ZDHHC11
GAPDH
1.0 1.7 1.9
1.0 1.7 2.0
C
MKN45
HA-ZDHHC11
CHX (h)
0 12 24 36 48
PD-L1
HA-ZDHHC11
GAPDH
Log(IOD time X/IOD time 0)
Time (h)
Vec
HA-ZDHHC11
D
AGS
MKN45
shZDHHC11
His-ubi
0 48 72 h
PD: Ni-NTA
IB: PD-L1
PD-L1
ZDHHC11
GAPDH
E
Tumor
DAPI
PD-L1
ZDHHC11
Merge
PBS
saEV
50 μm
Relative intensity
***
F
MKN45
YTN16
saEV
PD-L1
GAPDH
1.0 1.8 2.1 2.2
1.0 1.5 1.6 1.8
Relative mRNA expression of PD-L1
ns
PBS
saEV
G
MKN45
saEV
CHX (h)
0 6 12 24 48
PD-L1
GAPDH
Log(IOD time X/IOD time 0)
Time (h)
Ctrl
saEV
H
MKN45
YTN16
saEV
His-ubi
PD: Ni-NTA
IB: PD-L1
PD-L1
Vinculin

**Figure S3. *sa*EVs drive gastric cancer immune evasion by upregulating the palmitoyltransferase ZDHHC11 to stabilize PD-L1.**

(A) qRT-PCR analysis reveals unchanged ZDHHC3 and ZDHHC9 expression in GC cells following treatment with *sa*EV.

(B) Immunoblot analysis of PD-L1 upregulation in MKN1 and MKN45 cells following ZDHHC11 overexpression.

(C) Following ZDHHC11 overexpression, MKN45 cells were treated with cycloheximide (CHX; 100 μg/mL) for specified durations. PD-L1 protein stability was assessed by immunoblotting and quantified using ImageJ.

(D) Dox-inducible ZDHHC11-knockdown AGS and MKN45 cells were harvested following 6-hour treatment with 25 μM MG132. Cell lysates were subjected to nickel beads pulldown assays and immunoblotted with indicated antibodies.

(E) Immunofluorescence staining showing PD-L1 and ZDHHC11 upregulation in mice tumor tissues following oral administration of *sa*EVs (upper panel). The fluorescence intensities of PD-L1 and ZDHHC11 were quantified using ImageJ (lower panel).

(F) Immunoblot analysis showing PD-L1 protein upregulation after *sa*EV treatment (upper panel). qRT-PCR analysis reveals unchanged PD-L1 mRNA levels in *sa*EV-treated MKN45 and YTN16 cells (lower panel).

(G) Following *sa*EV treatment, MKN45 cells were treated with cycloheximide (CHX; 100 μg/ml) for specified durations. PD-L1 protein stability was assessed by immunoblotting and quantified using ImageJ.

(H) MKN45 and YTN16 cells treated with *sa*EV were harvested following 6-hour exposure to 25 μM MG132. Cell lysates were subjected to nickel beads pulldown assays and immunoblotted with indicated antibodies.

Data are presented as mean ± SD. *$p < 0.05$; **$p < 0.01$; ***$p < 0.001$; ns, not significant. EV, extracellular vesicle; CHX, cycloheximide; PD, pulldown; IB, immunoblot.

Figure S4

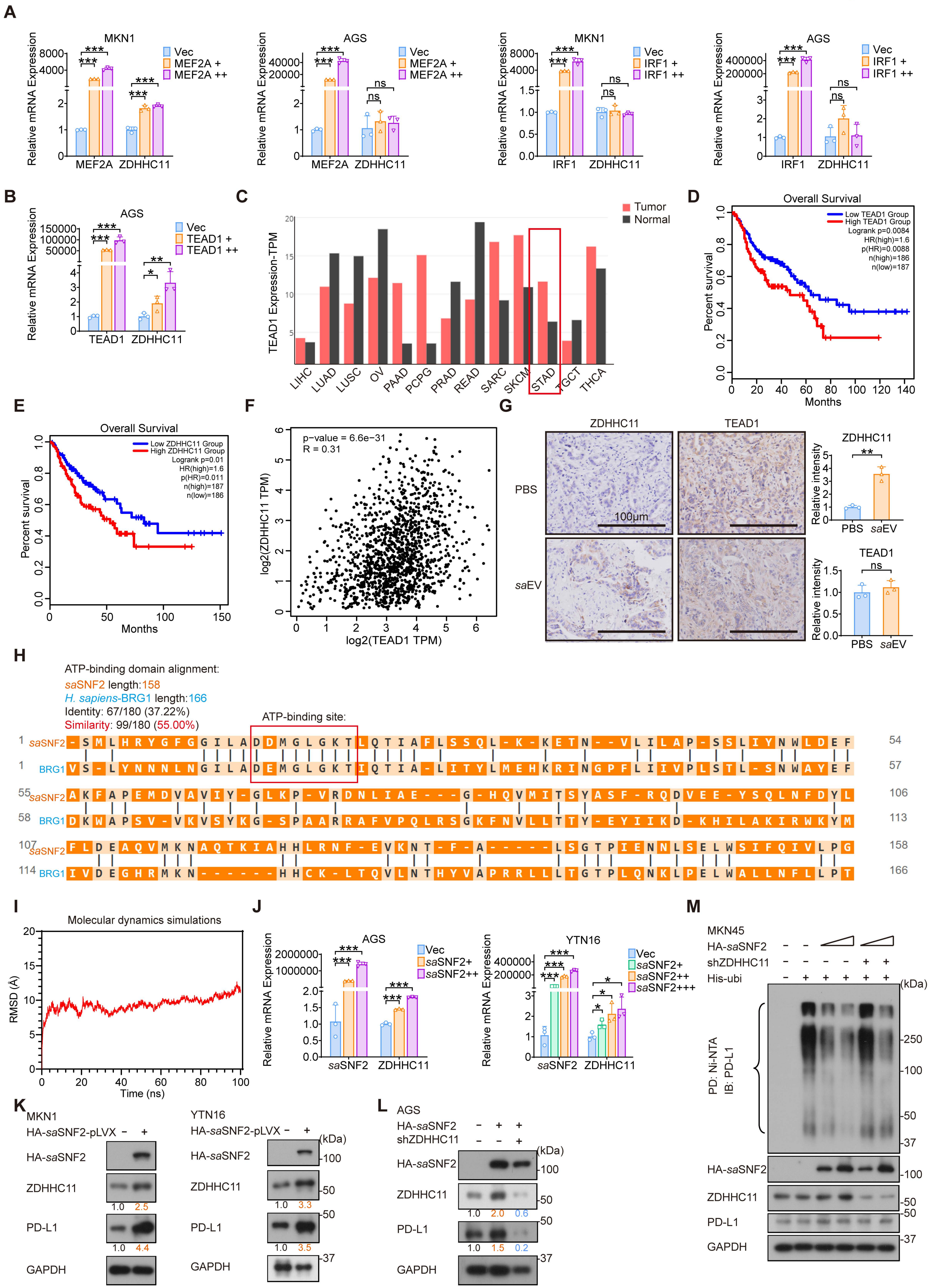

A
MKN1
AGS
Relative mRNA Expression
Vec
MEF2A +
MEF2A ++
IRF1 +
IRF1 ++
MEF2A
ZDHHC11
IRF1
***
**
*
ns
B
TEAD1 +
TEAD1 ++
TEAD1
C
Tumor
Normal
TEAD1 Expression-TPM
LIHC
LUAD
LUSC
OV
PAAD
PCPG
PRAD
READ
SARC
SKCM
STAD
TGCT
THCA
D
Overall Survival
Low TEAD1 Group
High TEAD1 Group
Logrank p=0.0084
HR(high)=1.6
p(HR)=0.0088
n(high)=186
n(low)=187
Percent survival
Months
E
Overall Survival
Low ZDHHC11 Group
High ZDHHC11 Group
Logrank p=0.01
HR(high)=1.6
p(HR)=0.011
n(high)=187
n(low)=186
Percent survival
Months
F
p-value = 6.6e−31
R = 0.31
log2(ZDHHC11 TPM)
log2(TEAD1 TPM)
G
ZDHHC11
TEAD1
PBS
saEV
100μm
Relative intensity
H
ATP-binding domain alignment:
saSNF2 length:158
H. sapiens-BRG1 length:166
Identity: 67/180 (37.22%)
Similarity: 99/180 (55.00%)
ATP-binding site:
saSNF2
BRG1
I
Molecular dynamics simulations
RMSD (Å)
Time (ns)
J
AGS
YTN16
Vec
saSNF2+
saSNF2++
saSNF2+++
Relative mRNA Expression
saSNF2
ZDHHC11
K
MKN1
YTN16
HA-saSNF2-pLVX
HA-saSNF2
ZDHHC11
PD-L1
GAPDH
(kDa)
1.0 2.5
1.0 4.4
1.0 3.3
1.0 3.5
L
AGS
HA-saSNF2
shZDHHC11
1.0 2.0 0.6
1.0 1.5 0.2
M
MKN45
HA-saSNF2
shZDHHC11
His-ubi
PD: Ni-NTA
IB: PD-L1
HA-saSNF2
ZDHHC11
PD-L1
GAPDH

**Figure S4. The *sa*EV-derived chromatin remodeler *sa*SNF2 enhances TEAD1-mediated transcription of the ZDHHC11 gene.**

(A and B) qRT-PCR analysis demonstrating the expression of ZDHHC11 in MKN1 and AGS cells following overexpression of MEF2A, IRF1 or TEAD1.

(C) Pan-cancer analysis via GEPIA revealed elevated TEAD1 expression in STAD tumor tissues compared to normal tissues, with differential expression patterns across other malignancies.

(D and E) Prognostic significance of TEAD1 and ZDHHC11 in gastrointestinal cancers (STAD/COAD/READ) was assessed using TCGA-derived Kaplan-Meier survival curves, with statistical significance determined by log-rank test (quartile-stratified).

(F) A positive correlation between TEAD1 and ZDHHC11 expression was observed in gastrointestinal cancers (STAD, COAD, READ, ESCA, LIHC) using TCGA data analyzed via GEPIA.

(G) Tumor tissue sections were immunohistochemically stained for ZDHHC11 and TEAD1 (left panel). Staining intensity was quantified using ImageJ and presented as bar graphs (right panel).

(H) Alignment of the ATP-binding domains in *sa*SNF2 and BRG1 (Homo sapiens) revealed a highly conserved ATP-binding site.

(I) Molecular dynamics simulations (100 ns) revealed the structural stability of the *sa*SNF2-TEAD1 complex, as quantified by root-mean-square deviation (RMSD) analysis.

(J) qRT-PCR analysis demonstrating an upregulation of ZDHHC11 in AGS and YTN16 cells following *sa*SNF2 overexpression.

(K and L) Immunoblot analysis of ZDHHC11 and PD-L1 expression in MKN1, YTN16 or AGS cells following *sa*SNF2 overexpression and ZDHHC11 knockdown (Dox-inducible).

(M) MKN45 cells carrying the indicated plasmids were harvested following 6-hour treatment with 25 μM MG132. Cell lysates were subjected to nickel beads pulldown assays and immunoblotted with indicated antibodies.

Data are presented as mean ± SD. $*p < 0.05$; $**p < 0.01$; $***p < 0.001$; ns, not significant. EV, extracellular vesicle; RMSD, root-mean-square deviation; STAD, Stomach adenocarcinoma; COAD, Colon adenocarcinoma; READ, Rectum adenocarcinoma; ESCA, Esophageal carcinoma; LIHC, Liver hepatocellular carcinoma; PD, pulldown; IB, immunoblot.

Figure S5

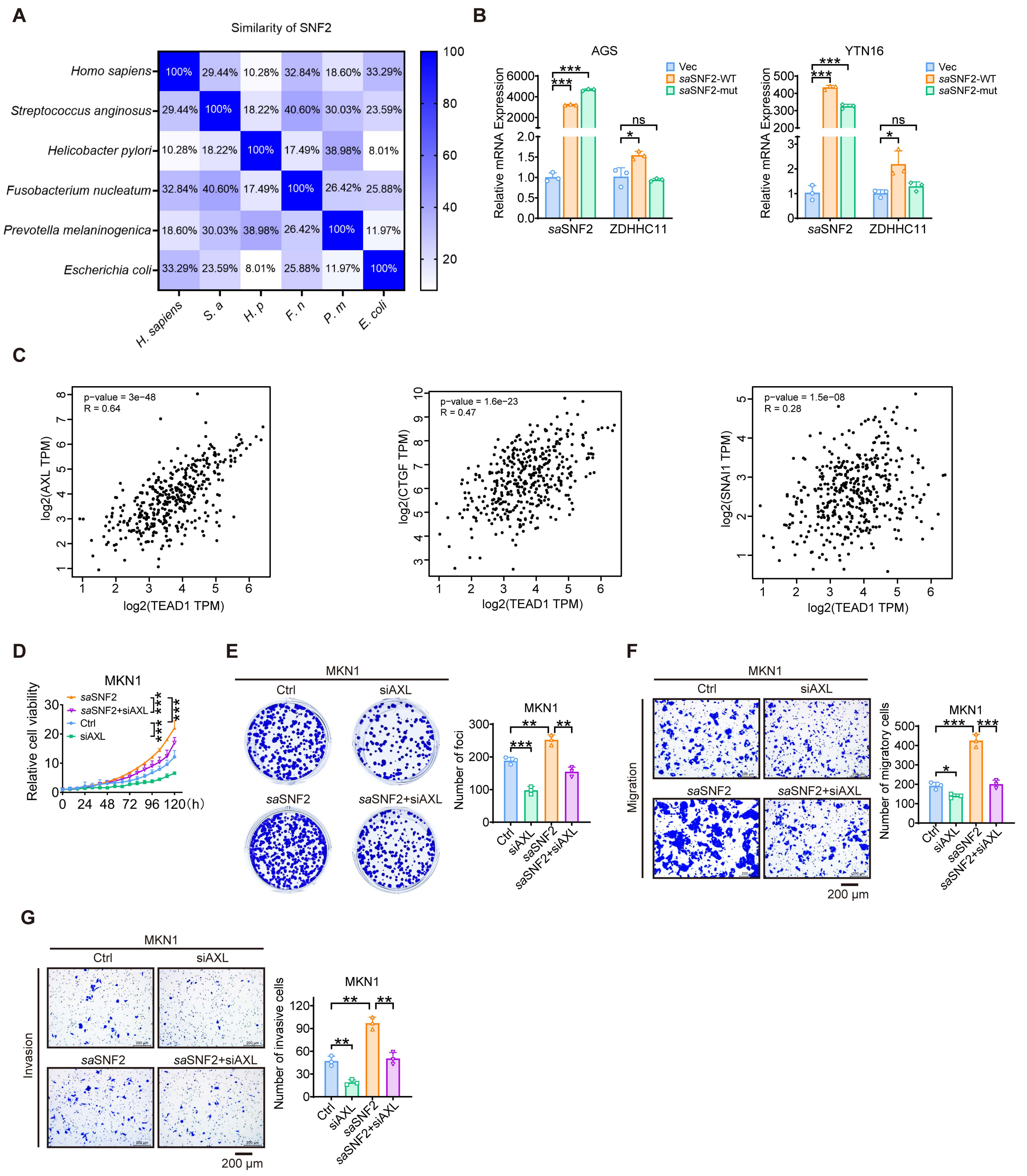
A
Similarity of SNF2
Homo sapiens
Streptococcus anginosus
Helicobacter pylori
Fusobacterium nucleatum
Prevotella melaninogenica
Escherichia coli
H. sapiens
S. a
H. p
F. n
P. m
E. coli
B
AGS
YTN16
Vec
saSNF2-WT
saSNF2-mut
Relative mRNA Expression
saSNF2
ZDHHC11
ns
C
p-value = 3e−48
R = 0.64
log2(AXL TPM)
log2(TEAD1 TPM)
p-value = 1.6e−23
R = 0.47
log2(CTGF TPM)
p-value = 1.5e−08
R = 0.28
log2(SNAI1 TPM)
D
MKN1
saSNF2
saSNF2+siAXL
Ctrl
siAXL
Relative cell viability
E
Ctrl
siAXL
saSNF2
saSNF2+siAXL
Number of foci
F
Migration
Number of migratory cells
200 μm
G
Invasion
Number of invasive cells
200 μm

**Figure S5. The ATPase motif of *sa*SNF2 is critical for TEAD1-mediated transcription and for assembly of the host BAF complex.**

(A) Comparative analysis of *sa*SNF2 protein sequence similarity in different species.

(B) qRT-PCR analysis demonstrating ZDHHC11 expression in AGS and YTN16 cells following overexpression of wild-type or mutant *sa*SNF2.

(C) Analysis of TCGA data using GEPIA revealed a positive correlation between TEAD1 and its target genes expression in stomach adenocarcinoma (STAD).

(D and E) The proliferative capacity and colony formation of MKN1 cells were examined after *sa*SNF2 overexpression and AXL knockdown (siRNA).

(F and G) The migratory and invasive capacities of MKN1 cells were assessed by Transwell assay after *sa*SNF2 overexpression and AXL knockdown (siRNA).

Data are presented as mean ± SD. *$p < 0.05$; **$p < 0.01$; ***$p < 0.001$; ns, not significant. STAD, Stomach adenocarcinoma.

### Figure S6

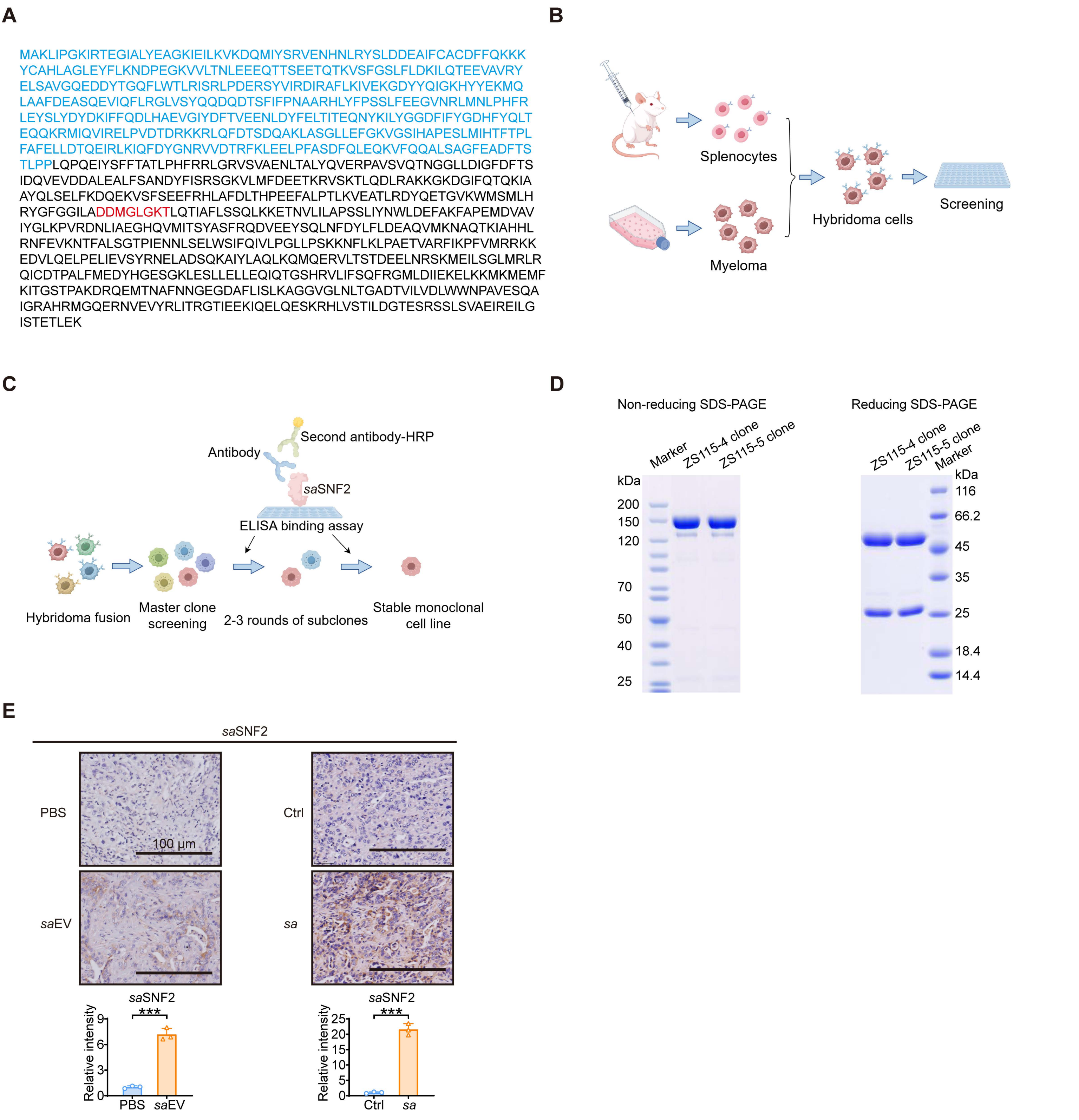

A
MAKLIPGKIRTEGIALYEAGKIEILKVKDQMIYSRVENHNLRYSLDDEAIFCACDFFQKKK
YCAHLAGLEYFLKNDPEGKVVLTNLEEEQTTSEETQTKVSFGSLFLDKILQTEEVAVRY
ELSAVGQEDDYTGQFLWTLRISRLPDERSYVIRDIRAFLKIVEKGDYYQIGKHYYEKMQ
LAAFDEASQEVIQFLRGLVSYQQDQDTSFIFPNAARHLYFPSSLFEEGVNRLMNLPHFR
LEYSLYDYDKIFFQDLHAEVGIYDFTVEENLDYFELTITEQNYKILYGGDFIFYGDHFYQLT
EQQKRMIQVIRELPVDTDRKKRLQFDTSDQAKLASGLLEFGKVGSIHAPESLMIHTFTPL
FAFELLDTQEIRLKIQFDYGNRVVDTRFKLEELPFASDFQLEQKVFQQALSAGFEADFTS
TLPPLQPQEIYSFFTATLPHFRRLGRVSVAENLTALYQVERPAVSVQTNGGLLDIGFDFTS
IDQVEVDDALEALFSANDYFISRSGKVLMFDEETKRVSKTLQDLRAKKGKDGIFQTQKIA
AYQLSELFKDQEKVSFSEEFRHLAFDLTHPEEFALPTLKVEATLRDYQETGVKWMSMLH
RYGFGGILADDMGLGKTLQTIAFLSSQLKKETNVLILAPSSLIYNWLDEFAKFAPEMDVAV
IYGLKPVRDNLIAEGHQVMITSYASFRQDVEEYSQLNFDYLFLDEAQVMKNAQTKIAHHL
RNFEVKNTFALSGTPIENNLSELWSIFQIVLPGLLPSKKNFLKLPAETVARFIKPFVMRRKK
EDVLQELPELIEVSYRNELADSQKAIYLAQLKQMQERVLTSTDEELNRSKMEILSGLMRLR
QICDTPALFMEDYHGESGKLESLLELLEQIQTGSHRVLIFSQFRGMLDIIEKELKKMKMEMF
KITGSTPAKDRQEMTNAFNNGEGDAFLISLKAGGVGLNLTGADTVILVDLWWNPAVESQA
IGRAHRMGQQERNVEVYRLITRGTIEEKIQELQESKRHLVSTILDGTESRSSLSVAEIREILG
ISTETLEK
B
Splenocytes
Myeloma
Hybridoma cells
Screening
C
Antibody
Second antibody-HRP
saSNF2
ELISA binding assay
Hybridoma fusion
Master clone screening
2-3 rounds of subclones
Stable monoclonal cell line
D
Non-reducing SDS-PAGE
Reducing SDS-PAGE
Marker
ZS115-4 clone
ZS115-5 clone
kDa
200
150
120
70
50
40
25
116
66.2
45
35
25
18.4
14.4
E
saSNF2
PBS
Ctrl
100 µm
saEV
sa
Relative intensity
***
0
3
6
9
5
10
15
20
25
PBS
saEV
Ctrl
sa

**Figure S6. Generation and validation of an anti-*sa*SNF2 monoclonal antibody.**

(A) Full-length amino acid sequence of *sa*SNF2. Blue highlights indicate the immunogen sequence used for antibody production. Red highlights mark the ATP-binding sites.

(B) Workflow for generating the *sa*SNF2-specific monoclonal antibody.

(C) Immunization schedule and serum titer assessment in Balb/c mice.

(D) Purity analysis of the final antibody under non-reducing (left panel) and reducing (right panel) conditions by SDS-PAGE.

(E) Tumor tissue sections were immunohistochemically stained for *sa*SNF2 expression (upper panel). Staining intensity was quantified using ImageJ and presented as bar graphs (lower panel).

### Table S1. Correlation of *sa*SNF2 expression with clinicopathological characteristics in 139 GC cases

| Variables | SNF2-low (*n* = 95) | SNF2-high (*n* = 44) | *P* value |
|---|---|---|---|
| Age (years) | 62 (25,84) | 62 (34,75) | 0.837 |
| Gender | | | |
| Male | 61 (64.2) | 35 (79.5) | 0.069 |
| Female | 34 (35.8) | 9 (20.5) | |
| CEA (ng/mL) | | | |
| <5 | 75 (78.9) | 33 (75.0) | 0.603 |
| ≥5 | 20 (21.1) | 11 (25.0) | |
| CA19-9 (U/mL) | | | |
| <37 | 78 (82.1) | 39 (88.6) | 0.326 |
| ≥37 | 17 (17.9) | 5 (11.4) | |
| CA125 (U/mL) | | | |
| <35 | 90 (94.7) | 41 (93.2) | 0.708 |
| ≥35 | 5 (5.3) | 3 (6.8) | |
| AFP (ng/mL) | | | |
| <8.78 | 91 (95.8) | 41 (93.2) | 0.679 |
| ≥8.78 | 4 (4.2) | 3 (6.8) | |
| T stage | | | |
| T1 | 16 (16.8) | 3 (6.7) | 0.012 |
| T2 | 13 (13.7) | 1 (2.3) | |
| T3 | 59 (62.1) | 31 (70.5) | |
| T4 | 7 (7.4) | 9 (20.5) | |
| N stage | | | |
| N0-N1 | 51 (53.7) | 15 (34.1) | 0.031 |
| N2-N3 | 44 (46.3) | 29 (65.9) | |
| M stage | | | |
| M0 | 83 (87.4) | 39 (88.6) | 0.832 |
| M1 | 12 (12.6) | 5 (11.4) | |
| TNM stage | | | |
| I-II | 52 (54.7) | 13 (29.5) | 0.006 |
| III-IV | 43 (45.3) | 31 (70.5) | |
| Differentiation | | | |
| Poor | 75 (78.9) | 28 (63.6) | 0.086 |
| Moderate | 17 (17.9) | 15 (34.1) | |
| well | 3 (3.2) | 1 (2.3) | |
| Vascular invasion | | | |
| Absence | 58 (61.1) | 29 (65.9) | 0.582 |
| Presence | 37 (38.9) | 15 (34.1) | |
| Perineural invasion | | | |
| Absence | 58 (61.1) | 26 (59.1) | 0.826 |
| Presence | 37 (38.9) | 18 (40.9) | |

The statistical analysis was performed using SPSS software. *P*-values were derived from two-sided chi-square tests, and significance was defined as $p < 0.05$. CEA, Carcinoembryonic antigen; CA19-9, Carbohydrate Antigen 19-9; CA125, Carbohydrate Antigen 125; AFP, α-fetoprotein.
